# PACS-1 variant protein is aberrantly localized in *C. elegans* model of PACS1/PACS2 syndromes

**DOI:** 10.1101/2024.04.22.590644

**Authors:** Dana T. Byrd, Ziyuan Christina Han, Christopher A. Piggott, Yishi Jin

## Abstract

PACS (Phosphofurin Acidic Cluster Sorting Protein) proteins are known for their roles in sorting cargo proteins to organelles and can physically interact with WD40 repeat-containing protein WDR37. PACS1, PACS2, and WDR37 variants are associated with multisystemic syndromes and neurodevelopmental disorders characterized by intellectual disability, seizures, developmental delays, craniofacial abnormalities, and autism spectrum disorder. However, the effects of syndromic variants on function *in vivo* remains unknown. Here, we report the expression pattern of *C. elegans* orthologs of PACS and WDR37 and their interaction. We show that cePACS-1 and ceWDR-37 co-localize to somatic cytoplasm of many types of cells, and are mutually required for expression, supporting a conclusion that the intermolecular dependence of PACS1/PACS2/PACS-1 and WDR37/WDR-37 is evolutionarily conserved. We further show that editing in PACS1 and PACS2 variants in cePACS-1 changes protein localization in multiple cell types, including neurons. Moreover, expression of human PACS1 can functionally complement *C. elegans* PACS-1 in neurons, demonstrating conserved functions of the PACS-WDR37 axis in an invertebrate model system. Our findings reveal effects of human variants and suggest potential strategies to identify regulatory network components that may contribute to understanding molecular underpinnings of PACS/WDR37 syndromes.

## Introduction

PACS (Phosphofurin Acidic Cluster Sorting Protein) family was first identified based on protein binding to a C-terminal acidic cluster motif within Furin protease (WAN *et al*. 1998) and is generally known to function in mediating cargo sorting or return to the trans-Golgi network (TGN) (YOUKER *et al*. 2009; THOMAS *et al*. 2017). PACS proteins are conserved from invertebrates to human (THOMAS *et al*. 2017). They are multi-domain proteins with N-terminus binding to Furin and C-terminal domains involved in autoregulation. Vertebrate genomes express two paralogs of PACS, and invertebrates generally have only one PACS homolog. Studies in multiple cell lines have revealed that interactions of acidic cluster motifs with the Furin Binding Region (FBR) of PACS1 and its paralogue, PACS2, can regulate membrane trafficking of multiple proteins, such as TRP, cyclic-nucleotide-gated channels, vesicular monoamine transporters and MHC-I (PIGUET *et al*. 2000; WAITES *et al*. 2001; KOTTGEN *et al*. 2005; JENKINS *et al*. 2009; YOUKER *et al*. 2009). Phosphorylation-mediated autoregulation of PACS proteins through the middle region (MR) also modifies their activities and subcellular site of action (SCOTT *et al*. 2003). PACS proteins have been shown to bind the conserved WD40 repeat-containing protein WDR37 in multiple protein interaction studies from invertebrates to mammals (LI *et al*. 2004; MALOVANNAYA *et al*. 2011; LIU *et al*. 2020; NAIR-GILL *et al*. 2021). In HEK293 cells, binding of WDR37 and PACS proteins appears to stabilize each other (NAIR-GILL *et al*. 2021). However, there is very limited *in vivo* analysis on endogenous PACS proteins, particularly in the nervous system.

Rapid advances in sequencing and data mining have hastened the discovery of genetic variants associated with neurodevelopmental disorders (NDD), intellectual disability (ID) and autism spectrum disorder (ASD). A major challenge remains to identify their genetic etiology. Recurrent *de novo* missense mutations in PACS1, PACS2, and WDR37 are recently reported to be associated with syndromes characterized by a similar set of symptoms. PACS1 syndrome was first reported in 2012 when an identical *de novo* variant in *pacs1* (c.607C>T; p.R203W) was found in two unrelated children with remarkably similar ID and facial features (SCHUURS-HOEIJMAKERS *et al*. 2012). In 2018, A PACS2 *de novo* mutation was reported in children with developmental and epileptic encephalopathy as well as facial dysmorphism similar to PACS1 syndrome (c.625G>A; p.E209K) (OLSON *et al*. 2018). In 2019, multiple variants of WDR37 were reported in children with a spectrum of symptoms including developmental delay, ID, and epilepsy (KANCA *et al*. 2019; REIS *et al*. 2019). Emerging experimental data and structural modeling analysis suggest that several WDR37 variants appear to cause unstable proteins (SOROKINA *et al*. 2021). It is unknown how PACS missense variants alter function and how all contribute to syndrome symptoms.

*C. elegans* has a single gene encoding PACS-1 and a single gene encoding WDR-37. In this study, we assessed endogenous localization of PACS-1 and WDR-37 by genomic insertion of fluorescent protein tags. We examined the effects of loss-of-function in *pacs-1* and *wdr-37*. We edited the endogenous *C. elegans pacs-1* gene to mimic human PACS syndrome-associated variants using CRISPR/Cas9. We show that PACS syndrome-associated variants alter PACS-1 localization. PACS-1 and WDR-37 are interdependent for expression and localization. Moreover, expression of human PACS1 can replace *C. elegans* PACS-1 function in neurons. Our work supports the evolutionary conservation of the PACS1-PACS2-WDR37 axis.

## Results and Discussion

### *C. elegans pacs-1* is broadly expressed in many tissues from embryogenesis to adult

*C. elegans* PACS-1 protein shows equal similarity to both human PACS1 (33% identical / 53% similar) and PACS2 (31% identical / 53% similar) (Figure S1a). Across phylogeny, the Furin Binding Region (FBR) is a highly conserved region of PACS proteins (YOUKER *et al*. 2009). The cePACS-1 FBR is 45% identical / 70% similar to hPACS-1 and 41% Identical / 67% Similar to hPACS-2 FBRs (Figure S1a). The residue altered in PACS1 syndrome, p.R203, lies within the FBR, corresponding to R116 in cePACS-1 (Figure S1b), whereas the position of the most common variant associated with PACS2 syndrome, p.E209, lies in the Middle Region (MR) and is conserved as E205 in cePACS-1 (ZANG *et al*. 2022) (Figure S1b).

To determine the endogenous expression and localization of PACS-1, we knocked in (KI) GFP or worm-optimized mScarlet (mSc) coding sequence into the *pacs-1* genomic locus using CRISPR/Cas9 (Figure S1d). As *pacs-1* is the first gene in an operon with *mdt-27* (Figure S1c), we verified GFP knock-in to *pacs-1* did not alter expression of *mdt-27* (Figure S1g)*. PACS-1::GFP(KI)* or *PACS-1::mSc (KI)* animals showed no overt phenotypes, including normal synaptic function tested by aldicarb sensitivity (Figure S1h,i). Expression of both KIs showed visible fluorescence in most cell types, most apparent in early embryos, germ cells, and neurons (Figure 1a-c). As mSc fluorescence quenches rapidly, we primarily focused on PACS-1::GFP in imaging analysis and verified key observations using PACS-1::mSc to eliminate potential bias of fluorescent proteins. PACS-1::GFP was first detected in early embryos and was enriched in the cytoplasm and near the plasma membrane of the AB cell in 2-cell embryos (Figure 1a). Throughout the early embryo cell divisions, expression remained higher in the somatic precursor cells and lower in the early germ cell precursor cells. In late larvae and adults, PACS-1::GFP expression was apparent in both the cytoplasm and near the membrane of germ cells (Figure 1b). In the head area, PACS-1::GFP was detectable in both the neuronal cell bodies adjacent to the pharynx and within the neuronal processes, or nerve ring (Figure 1c).

**Figure 1:**
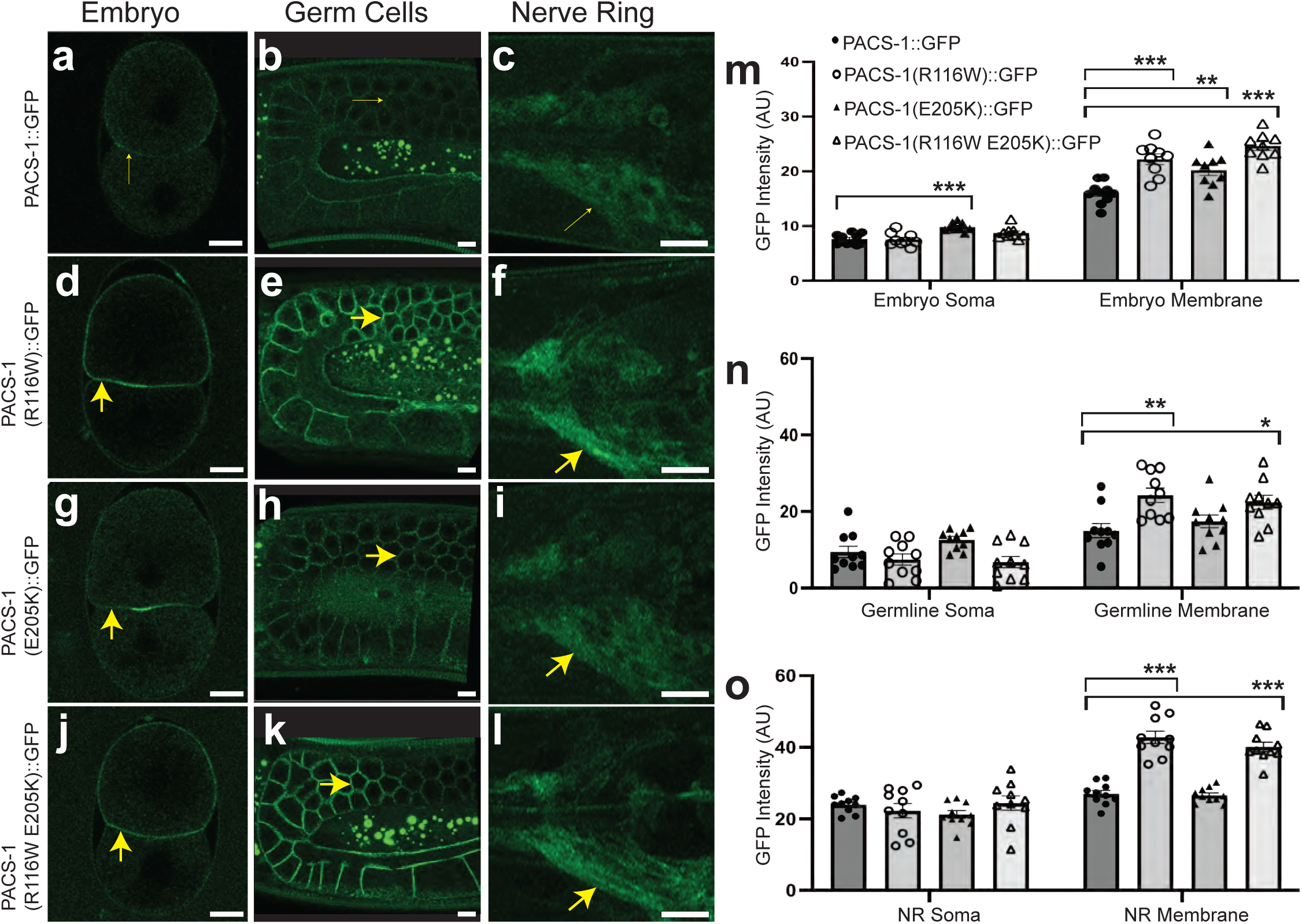
PACS1 and PACS2 disease variants alter localization of endogenously tagged *C. elegans* PACS-1 *in vivo*. (a-c) Expression of PACS-1::GFP in 2-cell embryos (a), meiotic germ cells and oocytes (b), and nerve ring (c). (d-f) Expression of PACS-1(R116W)::GFP in 2-cell embryos (d), meiotic germ cells and oocytes (e), and nerve ring (f). (g-i) Expression of PACS-1(E205K)::GFP in 2-cell embryos (g), meiotic germ cells and oocytes (h), and nerve ring (i). (j-l) Expression of PACS-1(R116W E205K)::GFP in 2-cell embryos (j), meiotic germ cells and oocytes (k), and nerve ring (l). (m-o) Quantification of wild type and variant PACS-1::GFP intensity in embryos (m), germ cells (n), and head neurons of nerve ring (NR) (o). Scale bars=1μm in a-b, d-e, g-h, and j-k; scale bars=10μm in c,f,i,l. Embryos oriented with anterior side on top. Adult animals oriented with anterior end to left. Imaged on Zeiss LSM800 Confocal microscope. 10-slice maximal projection for NR and 5-slice maximal projection for germ line with 0.5μm interval, single-slice images for embryos. Asterisks show results of one-way ANOVA with Tukey’s post-hoc test or Kruskal-Wallis H test followed by pairwise Wilcoxon rank sum exact test where *p≤0.05, **p≤0.01, ***p≤0.001. n≥10 for each genotype.

### *pacs-1* is not essential for development and growth, and human disease variants are also tolerated

To determine the function of *pacs-1*, we first analyzed a partial deletion mutation, *pacs-1*(*gk325*), shown as *pacs-1*(partial Δ) in Figure S1e, that removes the MR domain and the C-terminus. We observed grossly normal body morphology, size, growth, and movement. We also examined neuronal morphology and synapses using multiple neuronal markers and observed normal pattern in *pacs-1*(partial Δ) (Table S1). *pacs-1*(partial Δ) also did not significantly alter sensitivity to aldicarb or paraquat (Figures S1i and S2a-d), indicating that general synaptic transmission and stress response are likely intact. Additionally, during the process of gene-editing as described later, we obtained a *pacs-1* allele, designated as *pacs-1*(Δ) (Figure S1e), which has a premature stop after amino acid 29 as the result of a small deletion in the 5’ region of the gene and expressed greatly reduced *pacs-1* mRNA (Figure S1e, g). *pacs-1*(Δ) homozygous animals resembled *pacs-1(gk325)* in development, growth rate, and gross appearance of body shape and movement. Together, these observations indicate that despite early expression in embryos, *pacs-1* does not have essential roles in overall development.

We next edited *C. elegans* genomic DNA by CRISPR/Cas9 to generate animals expressing PACS-1(R116W) (Figure S1f) and examined the impact of the PACS1 human disease variant in a whole animal model. We found that like *pacs-1*(partial Δ), animals with *pacs-1*(R116W) showed no discernable differences in gross phenotypes such as body morphology, size, growth, or movement. It was previously reported that RNAi knockdown of *pacs-1* showed modest effect in the background of *dgk-1*/DGKQ diacylglycerol kinase *(lf)* (SIEBURTH *et al*. 2005). We constructed double mutants with *dgk-1(lf) and* observed that neither *pacs-1*(partial Δ) nor *pacs-1*(R116W) significantly modified aldicarb sensitivity (Figure S2c). We also tested whether *pacs-1* mutants could modify the behavior of *acr-2(gf),* which causes hyperactivation of an acetylcholine channel in the cholinergic motor neurons and exhibits seizure like behaviors (JOSPIN *et al*. 2009). We found neither *pacs-1*(partial Δ) nor *pacs-1*(R116W) affected *acr-2(gf)* behavior (Figure S2d).

PACS1 protein was first identified by its yeast 2-hybrid interaction with a psuedophosphorylated region of the Furin endoprotease (WAN *et al*. 1998). *C. elegans* KPC-1 and a related endoprotease, BLI-4, bear significant sequence similarity to the human Furin endoprotease, although the C-terminal regions do not align with the PACS1-interacting acidic cluster defined in human Furin (Q _768_ EECPSDSEEDEGRG) (WAN *et al*. 1998). Loss-of-function mutations in *kpc-1* cause a dramatic loss of PVD sensory neuron dendrites (SALZBERG *et al*. 2014)(Figure S2h). We found that neither *pacs-1*(partial 1−) nor *pacs-1*(R116W) significantly changed the dendritic structure of PVD neurons (Figure S2f-g). Like *kpc-1* single mutants, double mutants of *kpc-1* and *pacs-1* displayed a loss of PVD dendritic arborization at the same degree (Figure S2i-j). Given the high degree of conservation of the FBR, PACS proteins likely have conserved functions other than binding Furin.

### PACS-1 variants containing human PACS1 and PACS2 syndrome changes display increased accumulation near cell membrane

We next tested whether human PACS1 and PACS2 syndrome variants might affect expression or localization of *C. elegans* PACS-1 protein. We edited the variant amino acid position codons in PACS-1::GFP(KI) by CRISPR/Cas9 (Figure S1b and f). For PACS1 syndrome, cePACS-1 R116 was changed to W116 (human PACS1 R203W), and for PACS2 syndrome, cePACS-1 E205 was changed to K205 (human PACS2 E209K). While fluorescence intensity of PACS-1(R116W)::GFP and wild type PACS-1::GFP were similar in the soma of early embryos, oocytes, and head neurons, PACS-1(R116W)::GFP appeared significantly enriched in membrane surrounding early embryonic cells (Figure 1d and m), meiotic germ cells and oocytes (Figure 1e and n), as well as the neuronal processes of the nerve ring (Figure 1f and o). PACS-1(E205K)::GFP appeared slightly more enriched at the membrane of early embryonic cells (Figure 1g and m), but not statistically significantly enriched at membrane surrounding oocytes (Figure 1h and n) or in neuronal processes of the nerve ring (Figure 1i and o). It remains to be determined whether the differences between the levels of membrane enrichment of PACS-1(R116W)::GFP and PACS-1(E205K) reflect different impacts on protein function or different levels of impact on the same function.

We further tested how the PACS1 and PACS2 disease variants together affected PACS-1 protein expression. We made E205K as a second edit in animals expressing PACS-1(R116W)::GFP by CRISPR/Cas9 to generate PACS-1(R116W E205K)::GFP within the same protein (Figure S1f). PACS-1(R116W E205K)::GFP was enriched at membrane surrounding early embryonic cells (Figure 1j and m) and germ cells (Figure 1k and n) and in neuronal processes of the nerve ring (Figure 1l and o) to a similar extent as PACS-1(R116W)::GFP alone (Figure 1m-o). Thus, introducing the PACS2 variant in combination does not appear to change the effect of the PACS1 variant on localization.

### FBR domain of PACS-1 is important for protein stability

Since PACS1 and PACS2 disease-associated variants alter PACS-1 protein localization, we next sought to determine which domains are required for PACS-1 localization by generating endogenous in-frame deletions in PACS-1::GFP using CRISPR/Cas9. As mentioned above, *pacs-1* is the first gene in an operon with *mdt-27* (Figure S1c), which encodes a protein of the mediator family. Unexpectedly, we found that any genome editing that resulted in deletions in the 3’ region of *pacs-1*, corresponding to the coding sequences for the middle region (MR) and C-terminal region (CTR) of PACS-1, led to reduced expression of *mdt-27* mRNA. This observation may likely reflect genomic changes to the operon structure and confounds meaningful interpretation of observed changes in PACS-1 expression levels and localization patterns. We were able to obtain four independent in-frame deletions of the Furin binding region (FBR) denoted at *pacs-1*(1−FBR)::*gfp* (Figure S1e and Table S1) and confirmed expected mRNA sequences by sequencing cDNAs generated by RT-PCR from each line. In these mutants, PACS-1(1−FBR)::GFP showed diffuse localization with reduced levels of membrane association in early embryos (Figure 2d), germ cells (Figure 2e), and head neurons (Figure 2b and f). The reduced levels of PACS-1(1−FBR)::GFP are likely not due to changes in mRNA levels since levels of cDNA for both *pacs-1* and the downstream gene, *mdt-27* were not altered (Figure S1g). This analysis indicates that the FBR may stabilize the PACS-1 protein.

**Figure 2:**
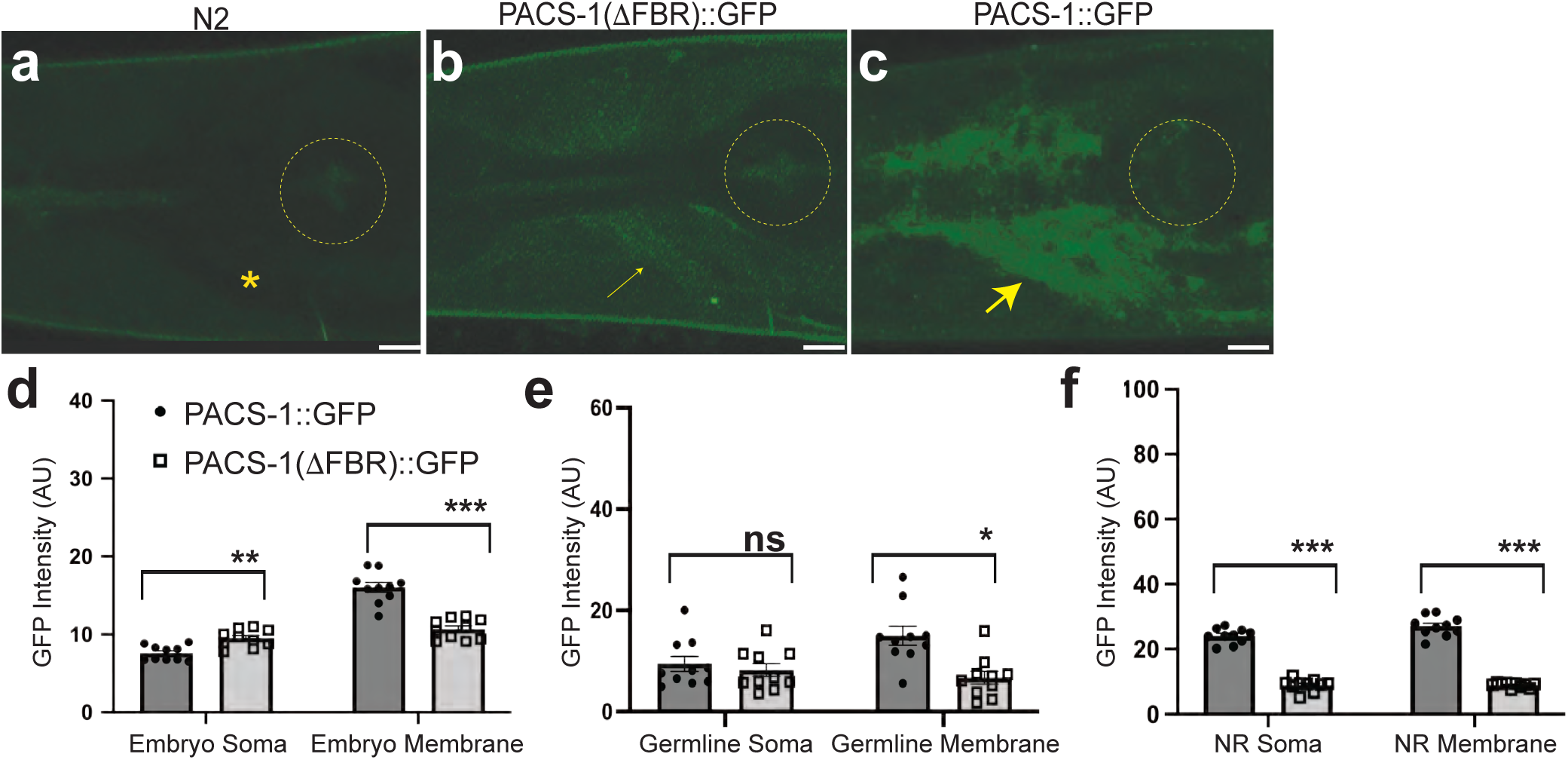
PACS-1 requires FBR domain for stability. (a-c) Head region of animals expressing no GFP (N2) (a), PACS-1(1−FBR)::GFP (b), and PACS-1::GFP (c). Adult animals oriented with anterior end to left. Scale bars =10μm. Yellow circles denote position of posterior pharyngeal bulb. (d-f) Quantification of PACS-1::GFP and PACS-1(1−FBR)::GFP expression in embryos (d), germ cells (e), and head neurons of nerve ring (NR) (f). Imaged on Zeiss LSM800 Confocal microscope. 10-slice maximal projection for NR with 0.5μm interval, single-slice images for germ line and embryos. Asterisks show results of one-way ANOVA with Tukey’s post-hoc test or Kruskal-Wallis H test followed by pairwise Wilcoxon rank sum exact test where *p≤0.05, **p≤0.01, ***p≤0.001, ns=not significant. n≥10 for each genotype.

### WDR-37 co-localizes with PACS-1 and is required for stable expression of PACS-1

WDR37/WDR-37 (Figure 4a) is a conserved interacting partner of PACS1/PACS-1 and PACS2 in human, mouse, and *C. elegans* (LI *et al*. 2004; NAIR-GILL *et al*. 2021; SOROKINA *et al*. 2021). *C. elegans* WDR-37 (C05D2.10a) and human WDR37 have similar structure of WD repeats (Figure 3a) and share 43% identity/59% similarity. To determine the function of *wdr-37*, we generated deletion alleles of *wdr-37, wdr-37(*1−), by CRISPR/Cas9 (Figure 3b). Like *pacs-1* loss-of-function, loss of *wdr-37* results in no overt phenotypes. *wdr-37(*1−) animals had normal growth rate, movement on plates, and neurotransmission measured by sensitivity to aldicarb. However, these animals showed a dramatic reduction of PACS-1::GFP in embryos (Figure 3l), germ cells (Figure 3e and m), and neurons (Figure 3f and n). Expressing wild type *wdr-37* cDNA in the neurons restored cytosolic PACS-1::GFP in *wdr-37(*1−) (Figure 3k). Reduced levels of PACS-1::GFP detected in *wdr-37(*1−) are most likely due to a dependence on WDR-37 for PACS-1 protein stability or reduced protein turnover since *pacs-1* mRNA/cDNA levels did not appear reduced *wdr-37(*1−) animals.

**Figure 3:**
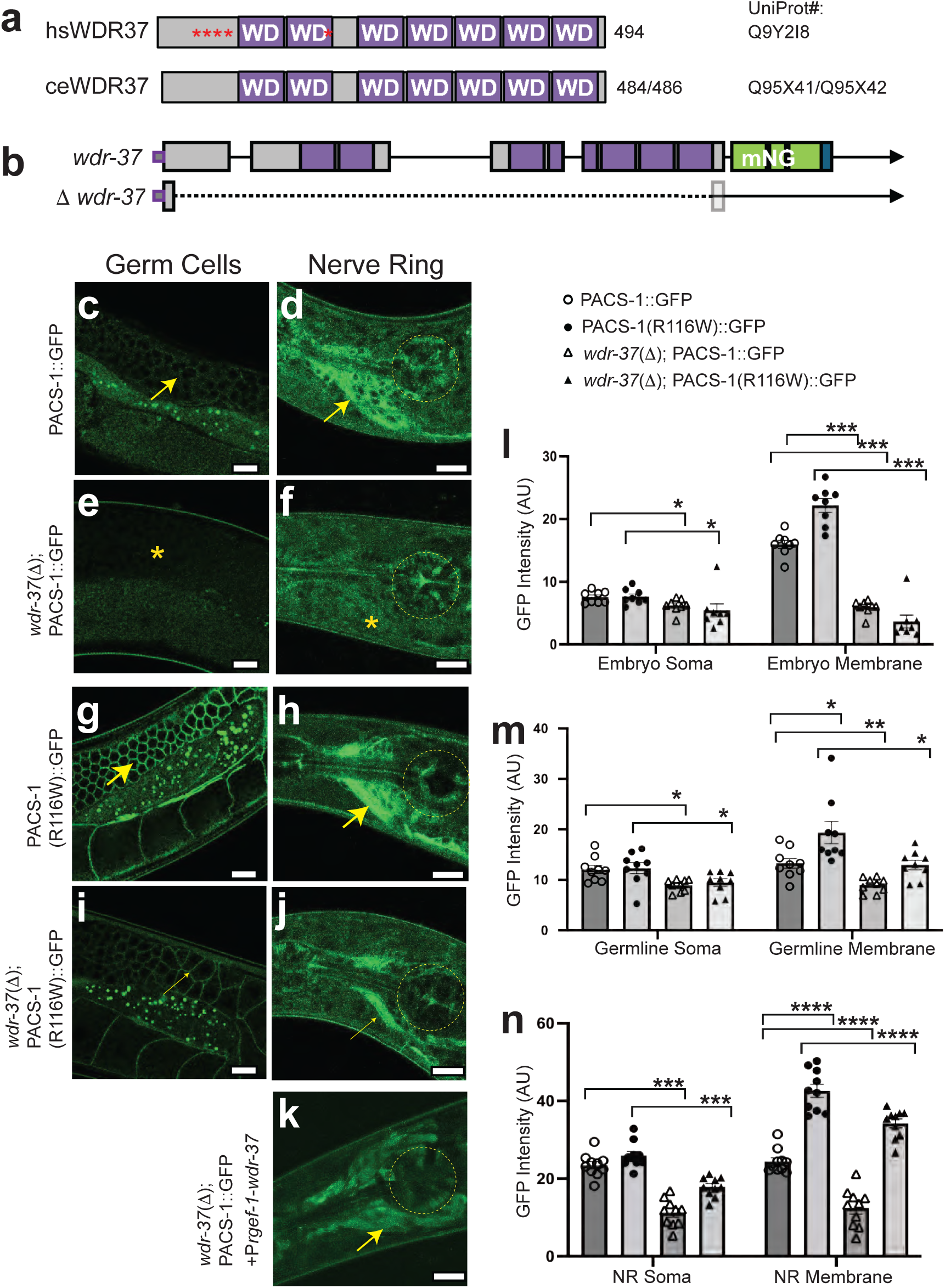
WDR-37 is required for PACS-1 localization in soma. (a) Schematic of human WDR37 and *C. elegans* WDR-37 proteins showing WD domain repeats (purple) and human variant positions (red asterisks). An alternative splice site at the 3’ end of the first intron of *C. elegans wdr-37* is predicted to generate a second isoform (C05D2.10b.1) that adds an additional 6 nucleotides, resulting in the 2 protein isoforms UniProt Q95X41 and Q95X42. However, in this study, all cDNAs sequenced matched the shorter C05D2.10a.1 isoform encoding UniProt Q95X41. (b) Gene structure of *wdr-37* (C05D2.10a.1) showing sites of CRISPR-generated genomic insertion of mNeonGreen (mNG) with Flag tag and deletion *wdr-37*(1−)(*ju1847*). (c-d) PACS-1::GFP in germ cells (c) and head neurons of nerve ring (d). (e-f) PACS-1::GFP in germ cells (e) and head neurons (f) of animals with *wdr-37* deletion (*wdr-37*(1−)). (g-h) Variant PACS-1(R116W)::GFP in germ cells (g) and head neurons of nerve ring (h). (i-j) Variant PACS-1(R116W)::GFP in germ cells (i) and head neurons (j) of animals with *wdr-37*(1−). (k) Expression of PACS-1::GFP in head neurons of *wdr-37*(1−) deletion animal rescued with expression of *Prgef-1*-*wdr-37* cDNA. 3/3 lines of animals showed rescued expression of PACS-1. 10-slice maximal projection with 0.5μm interval. (l-n) Quantification of PACS-1::GFP and PACS-1(R116W)::GFP expression with and without *wdr-37*(1−) in embryos (l), germ cells (m), and head neurons of nerve ring (NR) (n). Adult animals oriented with anterior end to left. Imaged on Zeiss LSM800 Confocal microscope. Yellow circles denote position of posterior pharyngeal bulb. Scale bars=10μm. Asterisks show results of one-way ANOVA with Tukey’s post-hoc test where *p≤0.05, **p≤0.01, ***p≤0.001, ****p≤0.0001, ns=not significant. n≥10 for each genotype.

**Figure 4:**
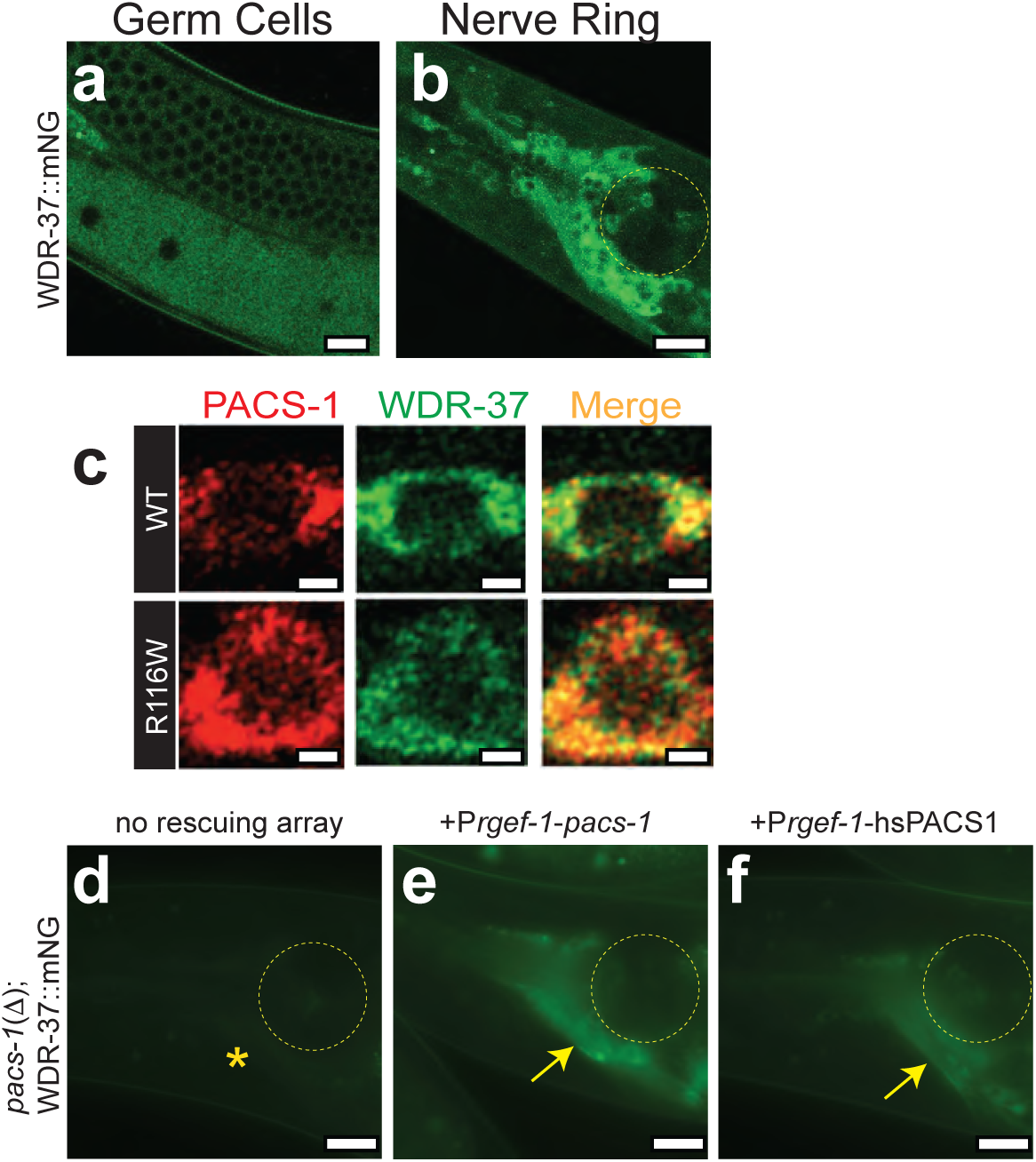
WDR-37 localization overlaps and depends on PACS-1, and PACS-1 can be functionally replaced by either *C. elegans* PACS-1 or human PACS1 in neurons. (a-b) WDR-37::mNG (*syb5283*) expression in meiotic germ cells and oocytes (a) and in head neurons (b). (c) Localization of PACS-1::mSc (top red) and variant PACS-1(R116W)::mSc (bottom red) overlap localization of WDR-37::mNG (green) as shown in merge (yellow). Imaged on Zeiss LSM900 airyscan microscope. (d-f) Expression of WDR-37::mNG in head neurons of *pacs-1*(1−) animals without rescuing extrachromosomal array (d), with rescuing neuronal expression of *C. elegans pacs-1* cDNA (*Prgef-1*-*pacs-1* cDNA) (e), and with rescuing neuronal expression of human PACS1 cDNA (*Prgef-1*-hsPACS1 cDNA) (f) 4/4 lines of animals expressing P*_rgef-1_*-*pacs-1* cDNA and 10/10 lines of animals expressing P*_rgef-1_*-hsPACS1 cDNA from extrachoromosomal arrays showed rescued levels of WDR-37::mNG expression in head neurons. Imaged on Zeiss Axio Imager M2 compound microscope under identical conditions. Adult animals oriented with anterior end to left. Yellow circles denote position of posterior pharyngeal bulb. Scale bars=10μm in a-b and d-f; scale bars=1μm in c.

We next tested whether variant PACS-1(R116W)::GFP is dependent on *wdr-37* for expression or localization. In *wdr-37(*1−) animals, we detected an overall reduction in levels of PACS-1(R116W)::GFP in embryos (Figure 3l), germ cells (Figure 3i and m), and neurons (Figure 3j and n); however, some PACS-1(R116W)::GFP remained enriched at the membranous regions between germ cells (Figure 3i and m) as well as the membranous nerve ring processes (Figure 4j and n). Therefore, *wdr-37* appears to be required for the stability of PACS-1 protein in the cytoplasm, whereas the stability of the membrane-associated pool of variant PACS-1(R116W) appears to be independent of WDR-37.

### PACS-1 regulates WDR-37 expression

To further examine how PACS-1 and WDR-37 might function together in *C. elegans*, we generated a C-terminal mNeonGreen (mNG)-tagged WDR-37 by CRISPR/Cas9 genomic insertion of mNG coding sequence in the endogenous *wdr-37* locus (Figure 3b). Consistent with their physical interaction, we found that WDR-37::mNG was expressed in a similar pattern as PACS-1::GFP, with most prominent expression in germ cells (Figure 4a) and neurons (Figure 4b). However, in germ cells, WDR-37::mNG was more prominent in the cytoplasm and not enriched at the surrounding membrane (Figure 4a). Within ventral motor neurons, PACS-1::mSc and WDR-37::mNG largely co-localized (Figure 4c top). PACS1 disease variant, PACS-1(R116W)::mSc, retained co-localization with WDR-37::mNG (Figure 4c bottom), suggesting that the PACS1 disease variant may not alter its physical interaction with WDR37.

Since murine PACS1 and WDR37 are mutually required for stability in HEK cells (NAIR-GILL *et al*. 2021), we tested whether *pacs-1* was required for expression of WDR-37::mNG and found that in *pacs-1*(1−) (Figure S1e), WDR-37::mNG levels were reduced in neurons of *pacs-1*(1−) animals (Figure 4d). Expression of *C. elegans pacs-1* cDNA in neurons restored neuronal expression of WDR-37::mNG in *pacs-1(*1−) animals (Figure 4e). Thus PACS-1 functions cell-autonomously to support stability or expression of WDR-37 in neurons.

### Expression of human PACS1 can complement *C. elegans* PACS-1 function in neurons

To test whether human PACS1 and *C. elegans* PACS-1 behave as functional orthologs, we expressed human PACS1 cDNA in *C. elegans* neurons and assessed whether it could also rescue the reduced levels of WDR-37::mNG observed in *pacs-1*(1−). We found that expression of human PACS1 cDNA in *C. elegans* neurons also rescued the neuronal expression of WDR-37::mNG (Figure 4f). Therefore, human PACS1 behaves as a functional ortholog of *C. elegans* PACS-1 and cell-autonomously supports stability or expression of WDR-37/WDR37 across phylogeny, further supporting *C. elegans* as a relevant model for understanding conserved function and identification of regulatory network components for the PACS1/PACS2/WDR37 axis.

### Conclusions

Pacs1/2 and Wdr37 are recently identified rare disease associated genes. In this study, we have reported the endogenous protein expression of PACS-1 and WDR-37 in living *C. elegans* and analyzed genetic null or strong loss of functions in each gene. We find that cePACS-1 and ceWDR-37 largely co-localize in somatic cytoplasm of many tissues and are mutually required for expression, providing *in vivo* evidence for the previous biochemical findings for mammalian PACS1 and WDR37 in cultured cells (NAIR-GILL *et al*. 2021). However, animals lacking *pacs-1* or *wdr-37* display normal development from embryo to adults. We have not detected discernable abnormal features in neuronal morphology, synapse development and neuron functions in multiple types of neurons tested. These data suggest that both genes may act in functionally redundant cellular processes. Through editing endogenous genes, we further characterized the effects of PACS1 and PACS2 syndromic variants and found that the human PACS1 and PACS2 variants in cePACS-1 altered protein localization. As studies of mammalian PACS proteins have revealed their broad roles in sorting and retrieval of multiple proteins, we speculate that PACS syndromic-associated proteins may disrupt cellular function by altering the subcellular distribution of a cohort or a select set of proteins depending on the types of cells. Lastly, we demonstrated that both *C. elegans* PACS-1 and human PACS1 can functionally complement PACS-1 in neurons. Our findings reveal the effects of human variants in a genetically amenable invertebrate model providing potential strategies to identify regulatory network components that may contribute to understanding molecular underpinnings of PACS/WDR37 syndromes.

## Acknowledgements

We thank our lab members for input throughout the project. We appreciate insightful discussions with J. Gleeson, T. Reddy, and members of the PACS1 Foundation. We are grateful for the seed fund from the PACS1 Foundation. Some strains were provided by the CGC, which is funded by NIH Office of Research Infrastructure Programs (P40 OD010440). This work was supported by R03NS121487(DTB and YJ) and NS R35NS127314(YJ).

## Author contributions

DTB and YJ conceptualized and acquired funding for this project. DTB, ZCH, and CAP collected and analyzed data. DTB, ZCH, and YJ wrote the paper. YJ supervised the project.

## Materials and methods

### Caenorhabditis elegans genetics

Wild type *C. elegans* is the N2 Bristol variant (BRENNER 1974). Strains were maintained under standard conditions on Nematode Growth Media (NGM) seeded with *E. coli* OP50 as described (BRENNER 1974), and listed in Table S1.

### CRISPR/Cas9 genome editing

For endogenous fluorescent tag insertions, *syb* knock-ins were purchased from SunyBiotech (Fuzhou, China). *pacs-1* deletion alleles were generated using two crRNAs: SD20246 and SD20670 for *pacs-1*(*ju2014*), shown as *pacs-1*(1−), and using an additional third crRNA: SD20144 for *pacs-1*(*ju1966*), shown as pacs-1(1−FBR) (Integrated DNA Technologies; Table S2: crRNA and primer sequences). The crRNAs were injected into *pacs-1*::*gfp*(*syb2274*) (SunyBiotech) with purified Cas9 (MacroLabs, University of California, Berkeley), trans-activating crRNA (tracrRNA; Integrated DNA Technologies) and *dpy-10* crRNA (TableS2), as described (PAIX *et al*. 2015). Dpy-10 F1 progeny were cultured individually, *pacs-1* DNA was amplified from F2 progeny with primers SD20247 and SD20230 (Table S2) to screen for deletions. Candidate deletions were confirmed by Sanger sequencing (Eton Bioscience, Inc.).

Human PACS variant alleles were generated by injecting N2, *pacs-1*::*gfp*(*syb2274*), and/or *pacs-1*::*mSc*(*syb2272*) (SunyBiotech) with SD20670 crRNA and SD21146 repair oligo for R116W edits or SD21142 crRNA and SD21147 repair oligo for E205K edits (Table S2) along with purified Cas9, tracrRNA, *unc-58* crRNA, and *unc-58*(*gf*) repair oligo as described (ARRIBERE *et al*. 2014). Unc-58 F1 progeny were cultured individually. *pacs-1* repair region was PCR amplified from F2 progeny with SD20406 and SD20407 for R116W or SD20229 and SD20230 for E205K and sequenced with SD20127 and SD20128 for R116W or SD20229 and SD20230 for E205K to detect and confirm variant edits (Table S2).

*wdr-37*(*ju1847*) deletion was generated by injecting N2 with three crRNAs: SD21143, SD21144, and SD21145 (Table S2) along with purified Cas9, tracrRNA, *unc-58* crRNA, and *unc-58*(*gf*) repair oligo as described (ARRIBERE *et al*. 2014). *wdr-37* deletions were screened and detected with SD20315, SD20316, and SD20317 (Table S2) in F2 progeny of each Unc-58 F1 animals.

All CRISPR-edited alleles were outcrossed at least twice with N2 and confirmed by genotyping (Table S2). Guide RNAs were selected using the CRISPR guide RNA selection tool (http://genome.sfu.ca/crispr/) and Integrated DNA Technologies crRNA design tools.

### Molecular biology and transgenesis

For RT-PCR, mixed stage worms were cultured under the same standard conditions. Total RNA was isolated using TRIzol (Thermo Fisher Scientific). All RNA samples were repurified using the TURBO DNA-free kit (Thermo Fisher Scientific). Reverse transcription was made using around 1μg of total RNA and the Superscript III RT kit (Thermo Fisher Scientific). To qualitatively assess the abundance of cDNA, all the cDNA samples were diluted into 200ng/uL and were amplified for 30x cycles using Primers SD20344 and SD20346 that target exon 6 and exon 8, respectively. SD20343 and SD20345 were used to confirm the sequence of cDNAs (Table S2). *mdt-27* in each cDNA sample was amplified using SD20382 and SD20383 (Table S2). *act-1* control was amplified from each sample using SD20368 and SD20369 (Table S2).

For *C. elegans* expression constructs, *pacs-1* and *wdr-37* cDNAs were amplified from an N2 cDNA library (PIGGOTT *et al*. 2021) with SD20414 and SD20415 for *pacs-1* or SD20416 and SD20417 for *wdr-37* (Table S2) using Phusion HF DNA Polymerase (NEB) and cloned into pCR8 by TOPO-TA cloning (Thermo Fisher Scientific). The *pacs-1* and *wdr-37* cDNAs were then recombined into Gateway plasmids containing the pan-neuronal promoter P*rgef-1* (pCZGY51 and pCZGY66) using Gateway LR Clonase II (Thermo Fisher Scientific), generating pCZGY3630 and pCZGY3615, respectively (Table S3). Human PACS1 cDNA (pENTR_PACS1_HA3; Gleeson Lab) was recombined into destination plasmid containing the pan-neuronal promoter P*rgef-1* (pCZGY66) using Gateway LR Clonase II (Thermo Fisher Scientific), generating pCZGY3631 (Table S3).

Transgenic lines were made by microinjection as previously described (MELLO *et al*. 1991). Plasmids and coinjection markers together with injection concentrations are listed in Table S3.

### Aldicarb sensitivity assay

To test aldicarb sensitivity, 30 day-1 adult animals were transferred to fresh plates containing 0.5 mM or 1 mM aldicarb. Animals were scored for paralysis every 15 or 30 min by gently touching the animal with a platinum wire. Final sample size for each assay was 20-25 animals due to some animals crawling off the plate.

### Paraquat sensitivity assay

Plates for measuring paraquat sensitivity were prepared by adding 18 µL of 1 M paraquat to 300 µL ddH_2_O, spread over seeded 5 cm NGM plates and dried 3 hours at room temperature. 20 day-1 adults were placed on each plate, incubated at 20°C, and monitored daily for survival.

### Fluorescence Microscopy

Animals were generally immobilized in a drop of M9 solution with 1 mM levamisole on a 4% agar pad or 10% agarose pad. Most fluorescence images were collected with a 63× oil immersion objective using a Zeiss LSM800 confocal microscope. Representative head neurons/nerve ring images are shown in 10-slice z-stack maximum projection at 0.5μm interval centered on middle of pharynx prepared using Fiji (ImageJ). Germ line images are presented as middle 5-slice z-stack maximum projection at 0.5μm interval. All embryo images are shown as single slice where the nucleus shows the biggest area. Soma and membrane regions were quantified by averaging the mean grey value from three regions of interest (ROIs). Images of ventral cord neurons expressing both WDR-37::mNG and PACS-1::mSc were collected with a 63× oil immersion objective using a Zeiss LSM900 airyscan microscope. For *pacs-1(ju2014)* rescue experiments, images of head neurons expressing WDR-37::mNG were collected with a 63× oil immersion objective on a Zeiss Axio Imager M2 compound microscope under identical settings.

### Statistical Analysis

Statistical analysis was performed using GraphPad Prism 9 or 10 (GraphPad Software, Inc.) and using RStudio (RStudio Team, 2020). Statistical significance was determined using one-way ANOVA with Tukey’s post-hoc test, non-parametric Kruskal-Wallis H test followed by pairwise Wilcoxon rank sum exact test, or unpaired, two-tailed *t*-tests. *P* > 0.05 was considered not significant (ns). *P* < 0.05 (*); *P* < 0.01 (**); *P* < 0.001 (***); and *P* < 0.0001 (****) were considered significant differences. Data are represented as mean ± SEM.

## Supplemental Material

**Figure S1:** *C. elegans pacs-1* encodes ortholog of human PACS1 and PACS2 with conserved disease variant positions.

**Figure S2:** Loss-of-function *pacs-1* and *pacs-1*(R116W) disease variant do not alter stress response, synaptic transmission, or dendritic morphology.

**Table S1:** Strains used in this study

**Table S2:** crRNA and oligo sequences

**Table S3:** DNA expression constructs

**Figure S1:**
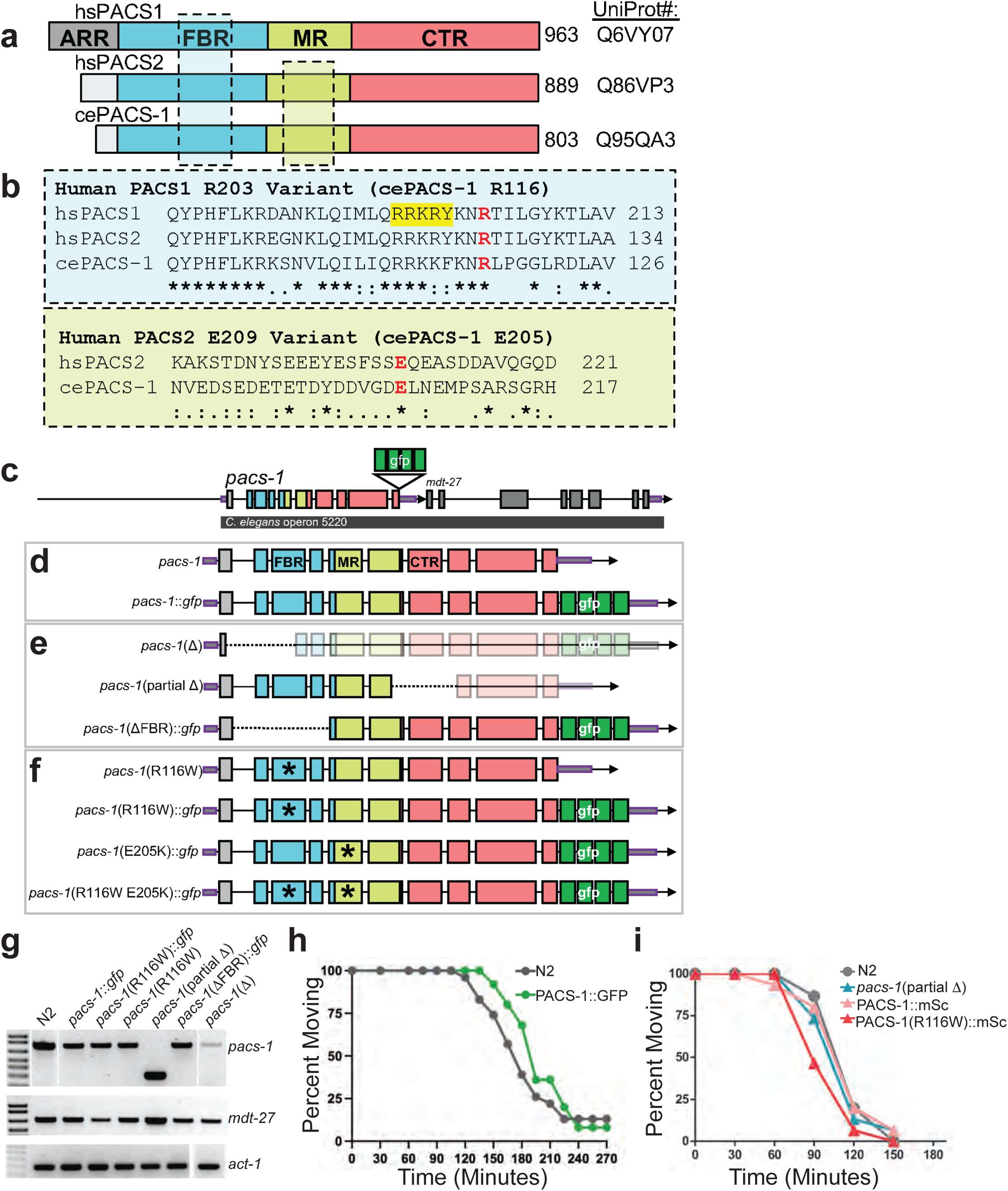
*C. elegans pacs-1* encodes ortholog of human PACS1 and PACS2 with conserved disease variant positions. (a) Human PACS1, PACS2, and *C. elegans* PACS-1 protein schematics. ARR is atrophin-1-related region (gray), FBR is furin binding region (cyan), MR is middle region (green), and CTR is C-terminal region (red). Boxed regions match alignments in B. (b) Clustal Omega sequence alignments of human disease variant regions: PACS1 R203W variant region (cyan box) and PACS2 E209K region (green box) with variant residues (red) and CK2 binding site (yellow highlight). (c) *C. elegans pacs-1* (T18H9.7) gene structure showing genomic insertion site of GFP (same site for mSc), and downstream gene within operon, *mdt-27*. (d) Wild type expected gene products for *pacs-1* without GFP (N2) and *pacs-1*::*gfp* (*pacs-1*(*syb2274*)). (e) Loss-of-function expected gene products from *pacs-1*(Δ) (*pacs-1*::*gfp*(*ju2014*)), *pacs-1*(partial Δ) (*pacs-1*(*gk325*)), and *pacs-1*(ΔFBR)::*gfp* (*pacs-1::gfp*(*ju1966*)). While *pacs-1*(Δ) was generated in the gfp KI strain, *pacs-1::gfp(syb2274),* coding region 3’ of the deletion, including gfp, is out of frame in *pacs-1*(Δ). (f) Human variant expected gene products *pacs-1*(R116W) (*pacs-1*(*ju1873*)), *pacs-1*(R116W)::*gfp* (*pacs-1::gfp* (*ju1823*)), *pacs-1*(E205K)::*gfp* (*pacs-1*::*gfp* (*ju1999*)), and *pacs-1*(R116W E205K)::*gfp* (*pacs-1*(R116W)::*gfp* (*ju2021*)). (g) rtPCR of *mdt-27*, *pacs-1* and *act-1* control using equivalent total cDNA from the following strains: N2, *pacs-1*::*gfp* (*syb2274*), *pacs-1*(R116W)::*gfp* (*ju1823*), *pacs-1*(R116W) (*ju1873*), *pacs-1*(partial Δ) (*gk325*), *pacs-1*(Δ FBR)::*gfp* (*ju1966*), and *pacs-1*(Δ) (*ju2014*). (h) Aldicarb sensitivity assay showing percentage of animals moving versus time in minutes for N2 and *pacs-1*::*gfp* (*syb2274*). GFP KI does not alter aldicarb sensitivity. Assayed on 0.5 mM Aldicarb plates. nζ20 for each genotype. (i) Aldicarb sensitivity assay showing percentage of animals moving versus time in minutes for N2, *pacs-1*(partial Δ) (*gk325*), *pacs-1*::*mSc* (*syb2272*), and *pacs-1*(R116W)::*mSc* (*ju1827*). PACS-1::mSC KI and *pacs-1*(partial Δ) loss of function do not alter aldicarb sensitivity. Assayed on 1 mM aldicarb plates. nζ20 for each genotype.

**Figure S2:**
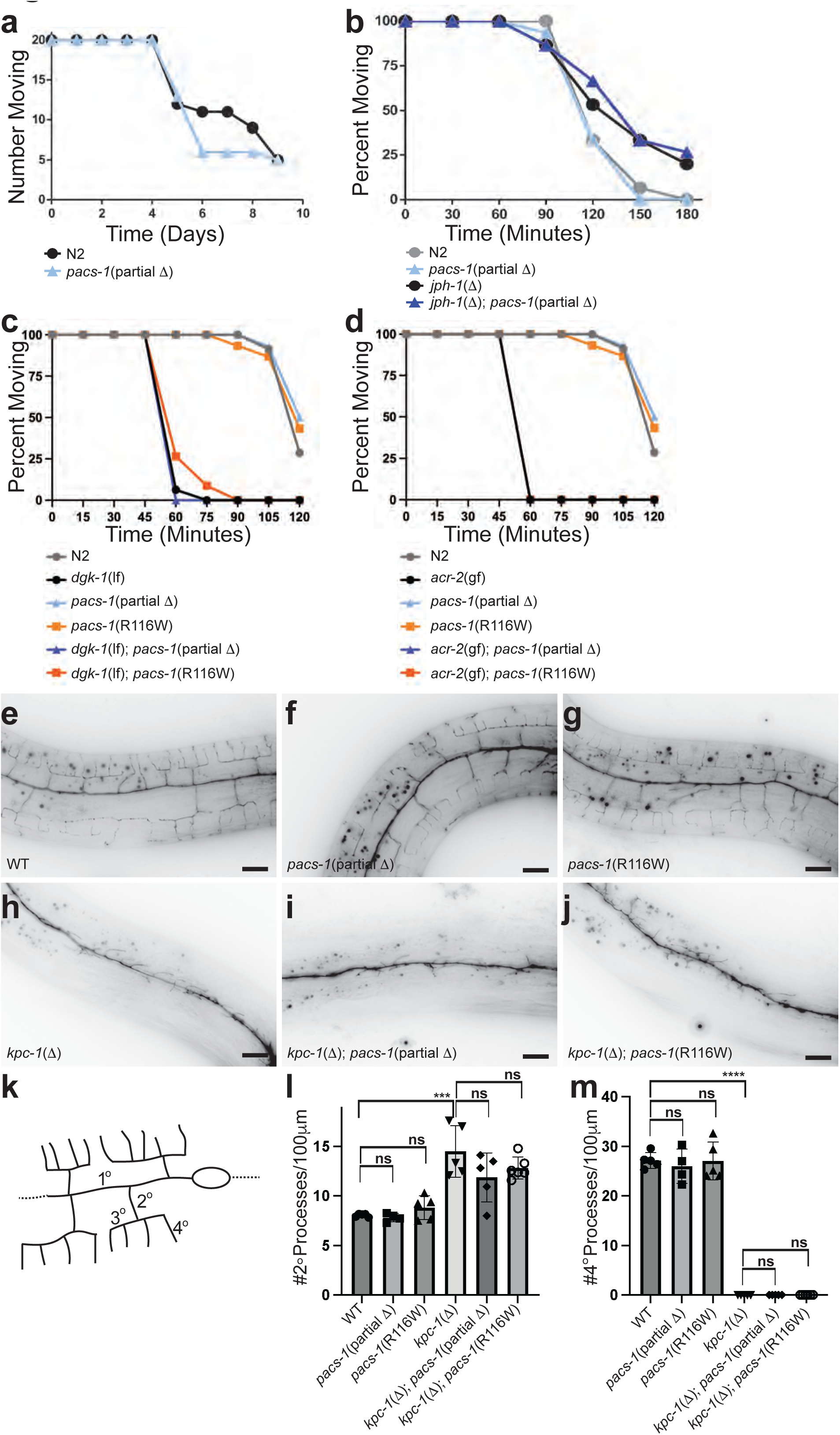
Loss-of-function in *pacs-1* and *pacs-1*(R116W) disease variant do not alter stress response, synaptic transmission, or dendritic morphology. (a) Paraquat sensitivity assay showing number of animals moving versus time in days. *pacs-1*(partial Δ) and wild type (N2) animals have similar sensitivity to paraquat. n=20 for each group. (b) Aldicarb sensitivity assay showing percentage of animals moving versus time in minutes. *pacs-1*(partial Δ) animals (light blue line) have similar aldicarb sensitivity as wild type (N2) animals (grey line) and do not significantly alter aldicarb resistance of *jph-1* animals (black line). Assayed on 1 mM aldicarb plates. n=20-30 for each genotype. (c) Aldicarb sensitivity assay showing percent of animals moving versus time in minutes. *pacs-1*(partial Δ) (*gk325* light blue line) and *pacs-1*(R116W) (*ju1873* light orange line) have similar aldicarb sensitivity as wild type animals (N2 gray line) and do not significantly alter aldicarb sensitivity of *dgk-1*(*lf*) (*nu62* black line). Assayed on 0.5 mM aldicarb plates. n=20-30 for each genotype. (d) Aldicarb sensitivity assay showing percent of animals moving versus time in minutes. *pacs-1*(partial Δ) (*gk325* blue line) and *pacs-1*(R116W) (*ju1873* orange line) do not significantly alter aldicarb sensitivity of *acr-2*(*gf*) (*n2420* black line). Assayed on 0.5 mM aldicarb plates. n=20-30 for each genotype. (e-j) Structure of PVD neurons visualized with *wyIs378* showing dendritic branches in wild type (e), *pacs-1*(R116W) (*ju1873*) (f), and *pacs-1*(partial Δ) (*gk325*) (g) animals, and reduced dendritic structure in *kpc-1*(Δ) (*gk8*) (h), *kpc-1*(Δ) (*gk8*); *pacs-1*(R116W) (*ju1873*) (i), and *kpc-1*(Δ) (*gk8*); *pacs-1*(partial Δ) (*gk325*) (j) animals. Images are composites from stack of 3 image planes taken on Zeiss AxioImager compound microscope. Animals oriented with anterior side to left. Scale bars=10μm. (k) Schematic of dendritic structure of PVD neuron with primary (1°), secondary (2°), tertiary (3°), and quaternary (4°) dendritic branches labeled. (l-m) Quantification of number of 2° branches (l) and 4° branches (m) per 100μM anterior of PVD cell body. Asterisks show results of unpaired t-tests where ***p≤0.001, ****p≤0.0001, ns=not significant. n≥4 for each genotype.

**Table S1:**
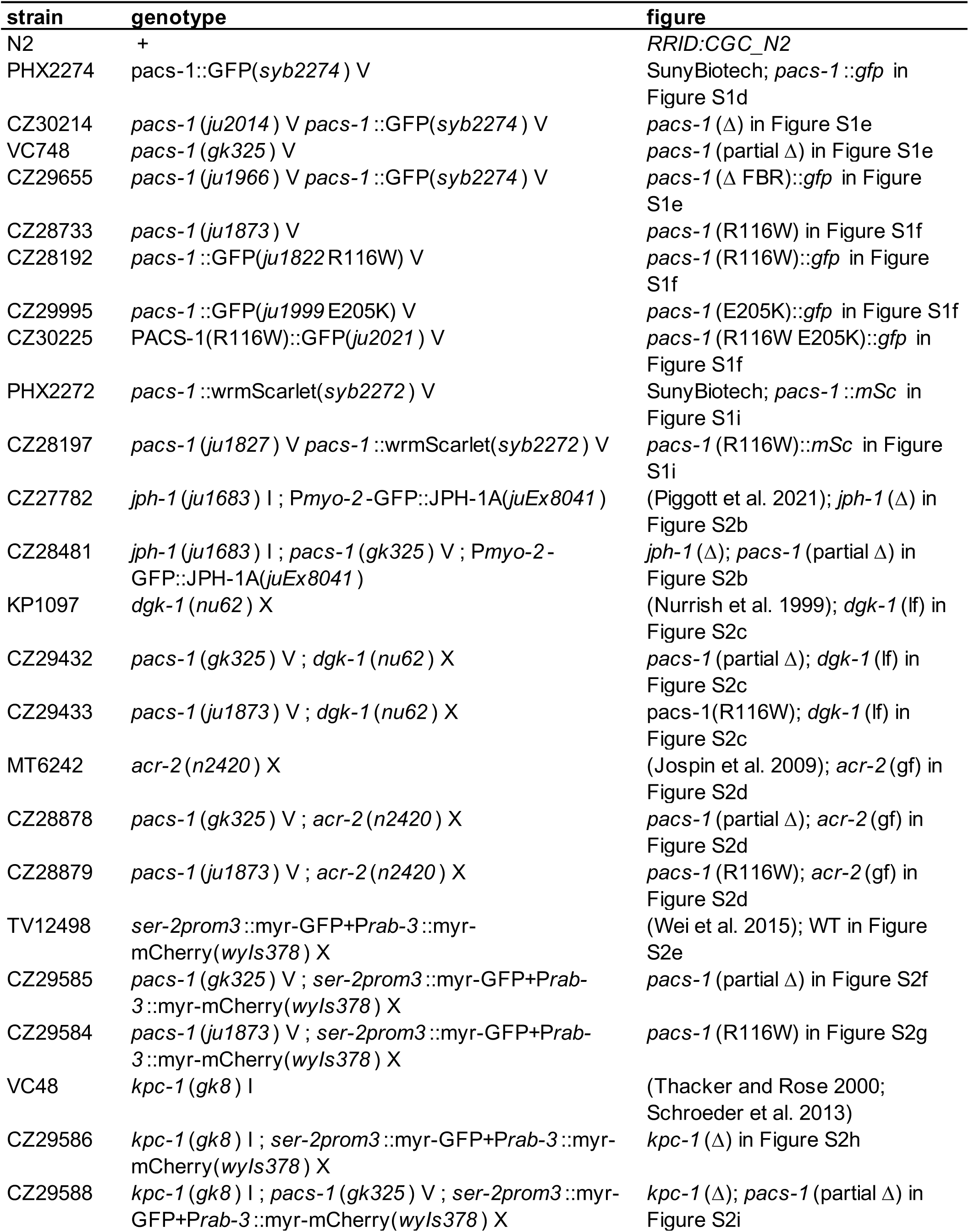
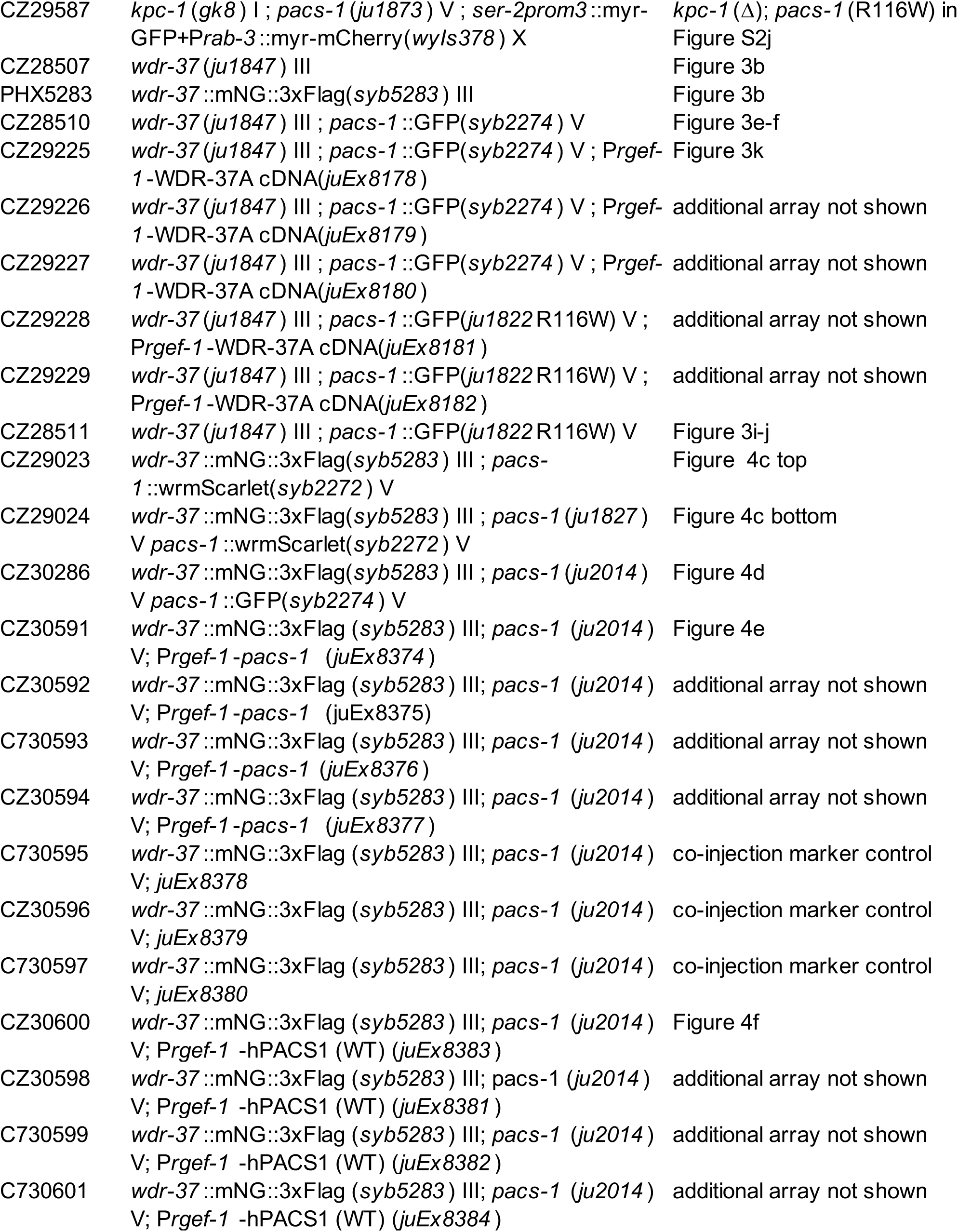
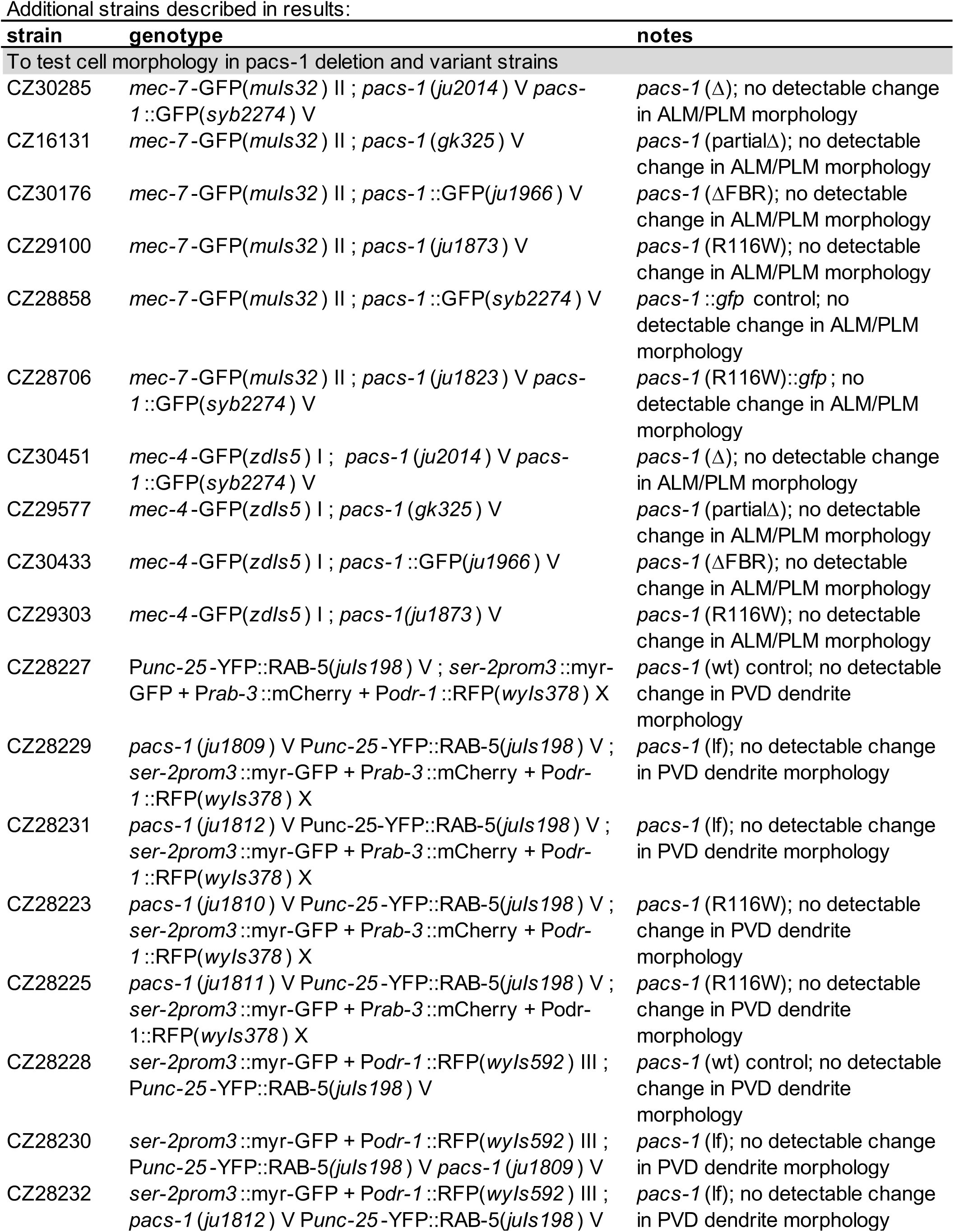
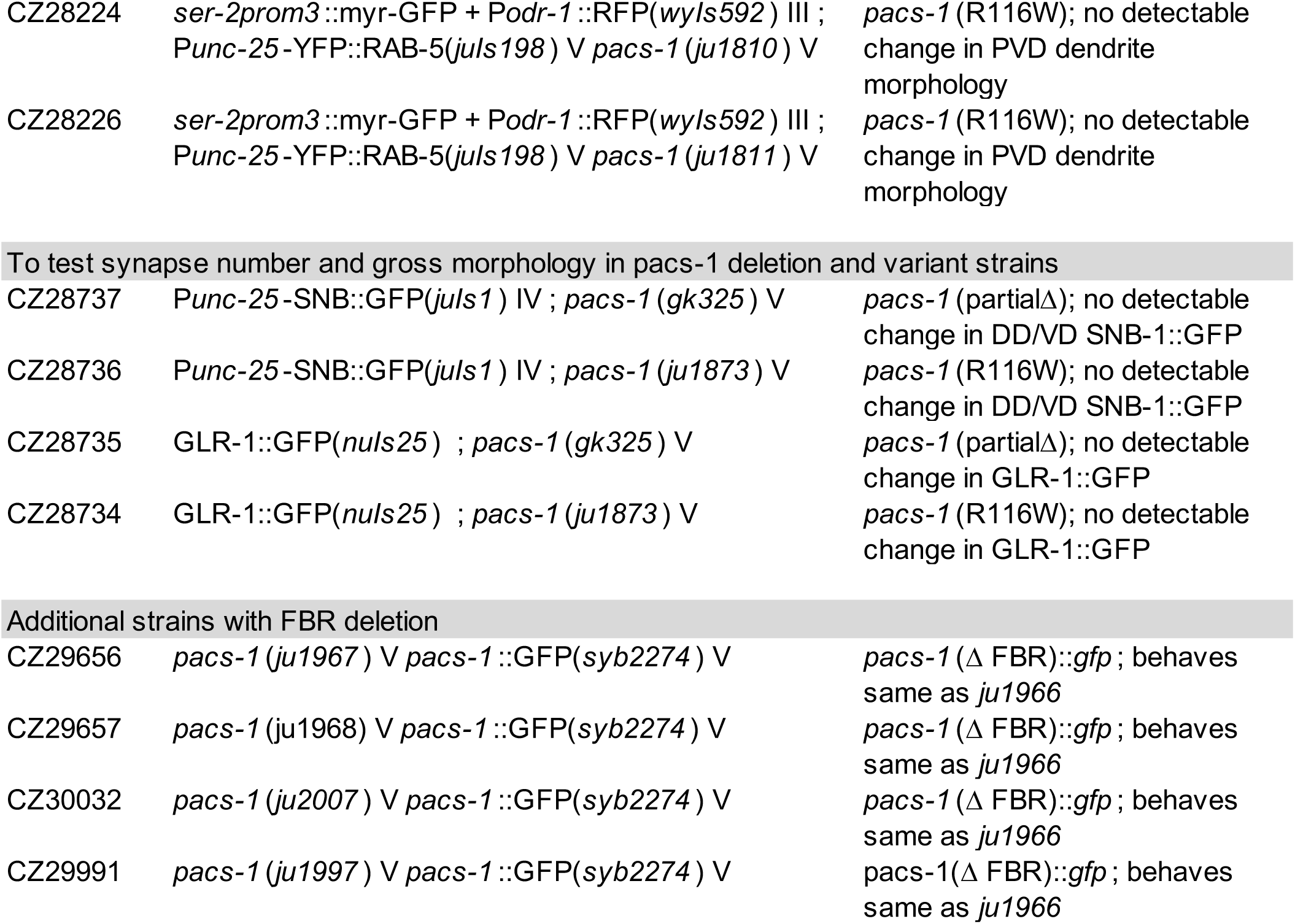
Strains used in this study.

| **strain** | **genotype** | **figure** |
| --- | --- | --- |
| N2 | + | *RRID:CGC_N2* |
| PHX2274 | *pacs-1::GFP(syb2274) V* | SunyBiotech; *pacs-1::gfp* in Figure S1d |
| CZ30214 | *pacs-1(ju2014) V pacs-1::GFP(syb2274) V* | *pacs-1(Δ)* in Figure S1e |
| VC748 | *pacs-1(gk325) V* | *pacs-1* (partial Δ) in Figure S1e |
| CZ29655 | *pacs-1(ju1966) V pacs-1::GFP(syb2274) V* | *pacs-1(Δ FBR)::gfp* in Figure S1e |
| CZ28733 | *pacs-1(ju1873) V* | *pacs-1* (R116W) in Figure S1f |
| CZ28192 | *pacs-1::GFP(ju1822 R116W) V* | *pacs-1* (R116W)::*gfp* in Figure S1f |
| CZ29995 | *pacs-1::GFP(ju1999 E205K) V* | *pacs-1* (E205K)::*gfp* in Figure S1f |
| CZ30225 | *PACS-1(R116W)::GFP(ju2021) V* | *pacs-1* (R116W E205K)::*gfp* in Figure S1f |
| PHX2272 | *pacs-1::wrmScarlet(syb2272) V* | SunyBiotech; *pacs-1::mSc* in Figure S1i |
| CZ28197 | *pacs-1(ju1827) V pacs-1::wrmScarlet(syb2272) V* | *pacs-1* (R116W)::*mSc* in Figure S1i |
| CZ27782 | *jph-1(ju1683) I ; Pmyo-2-GFP::JPH-1A(juEx8041)* | (Piggott et al. 2021); *jph-1(Δ)* in Figure S2b |
| CZ28481 | *jph-1(ju1683) I ; pacs-1(gk325) V ; Pmyo-2-GFP::JPH-1A(juEx8041)* | *jph-1(Δ)*; *pacs-1* (partial Δ) in Figure S2b |
| KP1097 | *dgk-1(nu62) X* | (Nurrish et al. 1999); *dgk-1* (lf) in Figure S2c |
| CZ29432 | *pacs-1(gk325) V ; dgk-1(nu62) X* | *pacs-1* (partial Δ); *dgk-1* (lf) in Figure S2c |
| CZ29433 | *pacs-1(ju1873) V ; dgk-1(nu62) X* | *pacs-1* (R116W); *dgk-1* (lf) in Figure S2c |
| MT6242 | *acr-2(n2420) X* | (Jospin et al. 2009); *acr-2* (gf) in Figure S2d |
| CZ28878 | *pacs-1(gk325) V ; acr-2(n2420) X* | *pacs-1* (partial Δ); *acr-2* (gf) in Figure S2d |
| CZ28879 | *pacs-1(ju1873) V ; acr-2(n2420) X* | *pacs-1* (R116W); *acr-2* (gf) in Figure S2d |
| TV12498 | *ser-2prom3::myr-GFP+Prab-3::myr-mCherry(wyls378) X* | (Wei et al. 2015); WT in Figure S2e |
| CZ29585 | *pacs-1(gk325) V ; ser-2prom3::myr-GFP+Prab-3::myr-mCherry(wyls378) X* | *pacs-1* (partial Δ) in Figure S2f |
| CZ29584 | *pacs-1(ju1873) V ; ser-2prom3::myr-GFP+Prab-3::myr-mCherry(wyls378) X* | *pacs-1* (R116W) in Figure S2g |
| VC48 | *kpc-1(gk8) I* | (Thacker and Rose 2000; Schroeder et al. 2013) |
| CZ29586 | *kpc-1(gk8) I ; ser-2prom3::myr-GFP+Prab-3::myr-mCherry(wyls378) X* | *kpc-1(Δ)* in Figure S2h |
| CZ29588 | *kpc-1(gk8) I ; pacs-1(gk325) V ; ser-2prom3::myr-GFP+Prab-3::myr-mCherry(wyls378) X* | *kpc-1(Δ)*; *pacs-1* (partial Δ) in Figure S2i |
| CZ29587 | *kpc-1(gk8)* I ; *pacs-1(ju1873)* V ; *ser-2prom3::myr-GFP+Prab-3::myr-mCherry(wyls378)* X | *kpc-1(Δ)*; *pacs-1* (R116W) in Figure S2j |
| CZ28507 | *wdr-37(ju1847)* III | Figure 3b |
| PHX5283 | *wdr-37::mNG::3xFlag(syb5283)* III | Figure 3b |
| CZ28510 | *wdr-37(ju1847)* III ; *pacs-1::GFP(syb2274)* V | Figure 3e-f |
| CZ29225 | *wdr-37(ju1847)* III ; *pacs-1::GFP(syb2274)* V ; *Prgef-1-WDR-37A cDNA(juEx8178)* | Figure 3k |
| CZ29226 | *wdr-37(ju1847)* III ; *pacs-1::GFP(syb2274)* V ; *Prgef-1-WDR-37A cDNA(juEx8179)* | additional array not shown |
| CZ29227 | *wdr-37(ju1847)* III ; *pacs-1::GFP(syb2274)* V ; *Prgef-1-WDR-37A cDNA(juEx8180)* | additional array not shown |
| CZ29228 | *wdr-37(ju1847)* III ; *pacs-1::GFP(ju1822R116W)* V ; *Prgef-1-WDR-37A cDNA(juEx8181)* | additional array not shown |
| CZ29229 | *wdr-37(ju1847)* III ; *pacs-1::GFP(ju1822R116W)* V ; *Prgef-1-WDR-37A cDNA(juEx8182)* | additional array not shown |
| CZ28511 | *wdr-37(ju1847)* III ; *pacs-1::GFP(ju1822R116W)* V | Figure 3i-j |
| CZ29023 | *wdr-37::mNG::3xFlag(syb5283)* III ; *pacs-1::wrmScarlet(syb2272)* V | Figure 4c top |
| CZ29024 | *wdr-37::mNG::3xFlag(syb5283)* III ; *pacs-1(ju1827)* V *pacs-1::wrmScarlet(syb2272)* V | Figure 4c bottom |
| CZ30286 | *wdr-37::mNG::3xFlag(syb5283)* III ; *pacs-1(ju2014)* V *pacs-1::GFP(syb2274)* V | Figure 4d |
| CZ30591 | *wdr-37::mNG::3xFlag(syb5283)* III; *pacs-1(ju2014)* V; *Prgef-1-pacs-1(juEx8374)* | Figure 4e |
| CZ30592 | *wdr-37::mNG::3xFlag(syb5283)* III; *pacs-1(ju2014)* V; *Prgef-1-pacs-1(juEx8375)* | additional array not shown |
| C730593 | *wdr-37::mNG::3xFlag(syb5283)* III; *pacs-1(ju2014)* V; *Prgef-1-pacs-1(juEx8376)* | additional array not shown |
| CZ30594 | *wdr-37::mNG::3xFlag(syb5283)* III; *pacs-1(ju2014)* V; *Prgef-1-pacs-1(juEx8377)* | additional array not shown |
| C730595 | *wdr-37::mNG::3xFlag(syb5283)* III; *pacs-1(ju2014)* V; *juEx8378* | co-injection marker control |
| CZ30596 | *wdr-37::mNG::3xFlag(syb5283)* III; *pacs-1(ju2014)* V; *juEx8379* | co-injection marker control |
| C730597 | *wdr-37::mNG::3xFlag(syb5283)* III; *pacs-1(ju2014)* V; *juEx8380* | co-injection marker control |
| CZ30600 | *wdr-37::mNG::3xFlag(syb5283)* III; *pacs-1(ju2014)* V; *Prgef-1-hPACS1(WT)(juEx8383)* | Figure 4f |
| CZ30598 | *wdr-37::mNG::3xFlag(syb5283)* III; *pacs-1(ju2014)* V; *Prgef-1-hPACS1(WT)(juEx8381)* | additional array not shown |
| C730599 | *wdr-37::mNG::3xFlag(syb5283)* III; *pacs-1(ju2014)* V; *Prgef-1-hPACS1(WT)(juEx8382)* | additional array not shown |
| C730601 | *wdr-37::mNG::3xFlag(syb5283)* III; *pacs-1(ju2014)* V; *Prgef-1-hPACS1(WT)(juEx8384)* | additional array not shown |

Additional strains described in results:
| strain | genotype | notes |
| --- | --- | --- |
| To test cell morphology in *pacs-1* deletion and variant strains |  |  |
| CZ30285 | *mec-7-GFP(muls32)* II ; *pacs-1(ju2014)* V *pacs-1::GFP(syb2274)* V | *pacs-1* ($\Delta$); no detectable change in ALM/PLM morphology |
| CZ16131 | *mec-7-GFP(muls32)* II ; *pacs-1(gk325)* V | *pacs-1* (partialΔ); no detectable change in ALM/PLM morphology |
| CZ30176 | *mec-7-GFP(muls32)* II ; *pacs-1::GFP(ju1966)* V | *pacs-1* (ΔFBR); no detectable change in ALM/PLM morphology |
| CZ29100 | *mec-7-GFP(muls32)* II ; *pacs-1(ju1873)* V | *pacs-1* (R116W); no detectable change in ALM/PLM morphology |
| CZ28858 | *mec-7-GFP(muls32)* II ; *pacs-1::GFP(syb2274)* V | *pacs-1::gfp* control; no detectable change in ALM/PLM morphology |
| CZ28706 | *mec-7-GFP(muls32)* II ; *pacs-1(ju1823)* V *pacs-1::GFP(syb2274)* V | *pacs-1* (R116W)::*gfp*; no detectable change in ALM/PLM morphology |
| CZ30451 | *mec-4-GFP(zdls5)* I ; *pacs-1(ju2014)* V *pacs-1::GFP(syb2274)* V | *pacs-1* ($\Delta$); no detectable change in ALM/PLM morphology |
| CZ29577 | *mec-4-GFP(zdls5)* I ; *pacs-1(gk325)* V | *pacs-1* (partialΔ); no detectable change in ALM/PLM morphology |
| CZ30433 | *mec-4-GFP(zdls5)* I ; *pacs-1::GFP(ju1966)* V | *pacs-1* (ΔFBR); no detectable change in ALM/PLM morphology |
| CZ29303 | *mec-4-GFP(zdls5)* I ; *pacs-1(ju1873)* V | *pacs-1* (R116W); no detectable change in ALM/PLM morphology |
| CZ28227 | *Punc-25-YFP::RAB-5(juls198)* V ; *ser-2prom3::myr-GFP + Prab-3::mCherry + Podr-1::RFP(wyls378)* X | *pacs-1* (wt) control; no detectable change in PVD dendrite morphology |
| CZ28229 | *pacs-1(ju1809)* V *Punc-25-YFP::RAB-5(juls198)* V ; *ser-2prom3::myr-GFP + Prab-3::mCherry + Podr-1::RFP(wyls378)* X | *pacs-1* (lf); no detectable change in PVD dendrite morphology |
| CZ28231 | *pacs-1(ju1812)* V *Punc-25-YFP::RAB-5(juls198)* V ; *ser-2prom3::myr-GFP + Prab-3::mCherry + Podr-1::RFP(wyls378)* X | *pacs-1* (lf); no detectable change in PVD dendrite morphology |
| CZ28223 | *pacs-1(ju1810)* V *Punc-25-YFP::RAB-5(juls198)* V ; *ser-2prom3::myr-GFP + Prab-3::mCherry + Podr-1::RFP(wyls378)* X | *pacs-1* (R116W); no detectable change in PVD dendrite morphology |
| CZ28225 | *pacs-1(ju1811)* V *Punc-25-YFP::RAB-5(juls198)* V ; *ser-2prom3::myr-GFP + Prab-3::mCherry + Podr-1::RFP(wyls378)* X | *pacs-1* (R116W); no detectable change in PVD dendrite morphology |
| CZ28228 | *ser-2prom3::myr-GFP + Podr-1::RFP(wyls592)* III ; *Punc-25-YFP::RAB-5(juls198)* V | *pacs-1* (wt) control; no detectable change in PVD dendrite morphology |
| CZ28230 | *ser-2prom3::myr-GFP + Podr-1::RFP(wyls592)* III ; *Punc-25-YFP::RAB-5(juls198)* V *pacs-1(ju1809)* V | *pacs-1* (lf); no detectable change in PVD dendrite morphology |
| CZ28232 | *ser-2prom3::myr-GFP + Podr-1::RFP(wyls592)* III ; *pacs-1(ju1812)* V *Punc-25-YFP::RAB-5(juls198)* V | *pacs-1* (lf); no detectable change in PVD dendrite morphology |
| CZ28224 | *ser-2prom3::myr-GFP + Podr-1::RFP(wyls592) III ; Punc-25-YFP::RAB-5(juls198) V pacs-1 (ju1810) V* | *pacs-1* (R116W); no detectable change in PVD dendrite morphology |
| CZ28226 | *ser-2prom3::myr-GFP + Podr-1::RFP(wyls592) III ; Punc-25-YFP::RAB-5(juls198) V pacs-1 (ju1811) V* | *pacs-1* (R116W); no detectable change in PVD dendrite morphology |
| To test synapse number and gross morphology in *pacs-1* deletion and variant strains |  |  |
| CZ28737 | *Punc-25-SNB::GFP(juls1) IV ; pacs-1 (gk325) V* | *pacs-1* (partialΔ); no detectable change in DD/VD SNB-1::GFP |
| CZ28736 | *Punc-25-SNB::GFP(juls1) IV ; pacs-1 (ju1873) V* | *pacs-1* (R116W); no detectable change in DD/VD SNB-1::GFP |
| CZ28735 | *GLR-1::GFP(nuls25) ; pacs-1 (gk325) V* | *pacs-1* (partialΔ); no detectable change in GLR-1::GFP |
| CZ28734 | *GLR-1::GFP(nuls25) ; pacs-1 (ju1873) V* | *pacs-1* (R116W); no detectable change in GLR-1::GFP |
| Additional strains with FBR deletion |  |  |
| CZ29656 | *pacs-1 (ju1967) V pacs-1::GFP(syb2274) V* | *pacs-1* ($\Delta$ FBR)::*gfp* ; behaves same as *ju1966* |
| CZ29657 | *pacs-1 (ju1968) V pacs-1::GFP(syb2274) V* | *pacs-1* ($\Delta$ FBR)::*gfp* ; behaves same as *ju1966* |
| CZ30032 | *pacs-1 (ju2007) V pacs-1::GFP(syb2274) V* | *pacs-1* ($\Delta$ FBR)::*gfp* ; behaves same as *ju1966* |
| CZ29991 | *pacs-1 (ju1997) V pacs-1::GFP(syb2274) V* | *pacs-1* ($\Delta$ FBR)::*gfp* ; behaves same as *ju1966* |

**Table S2.**
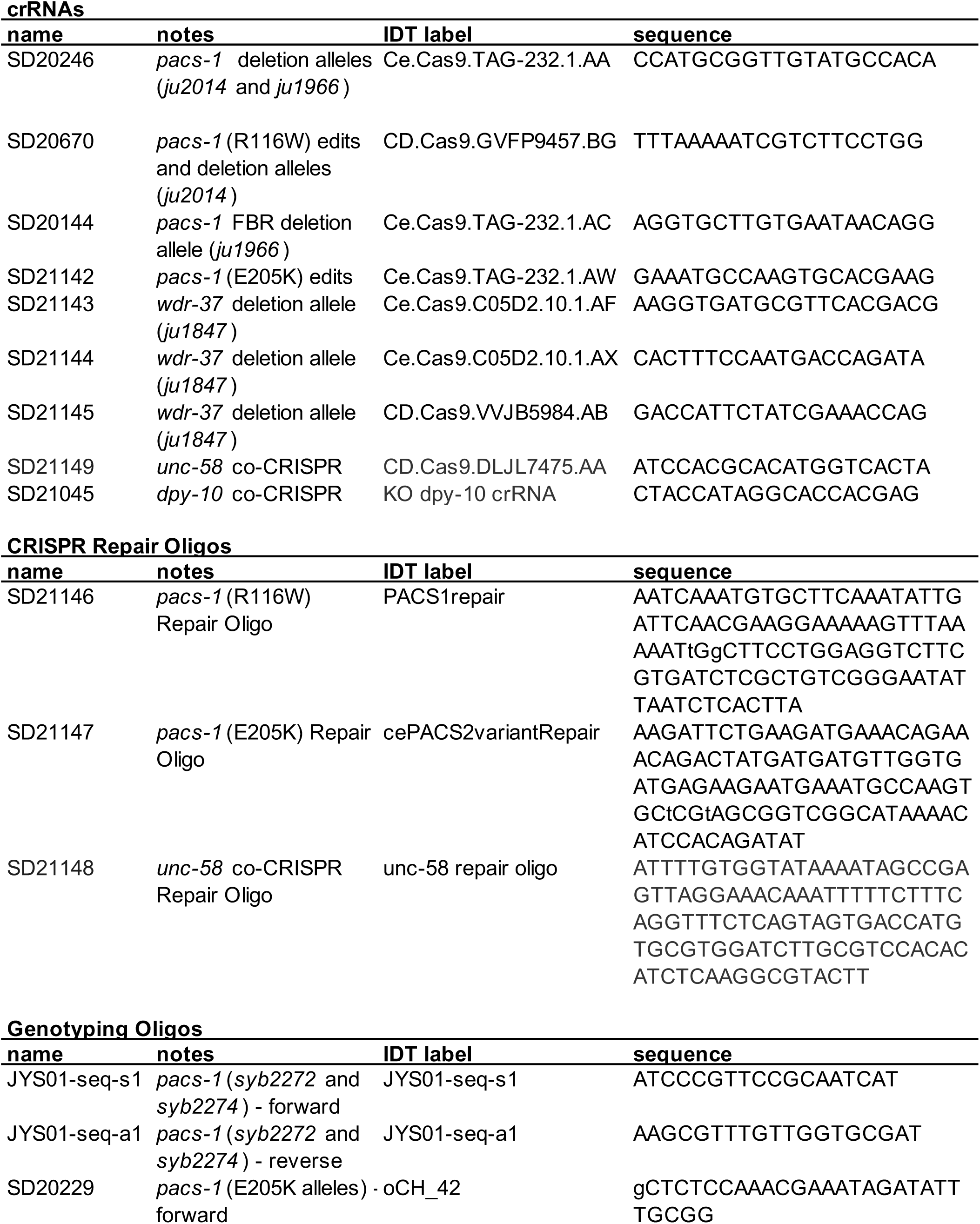
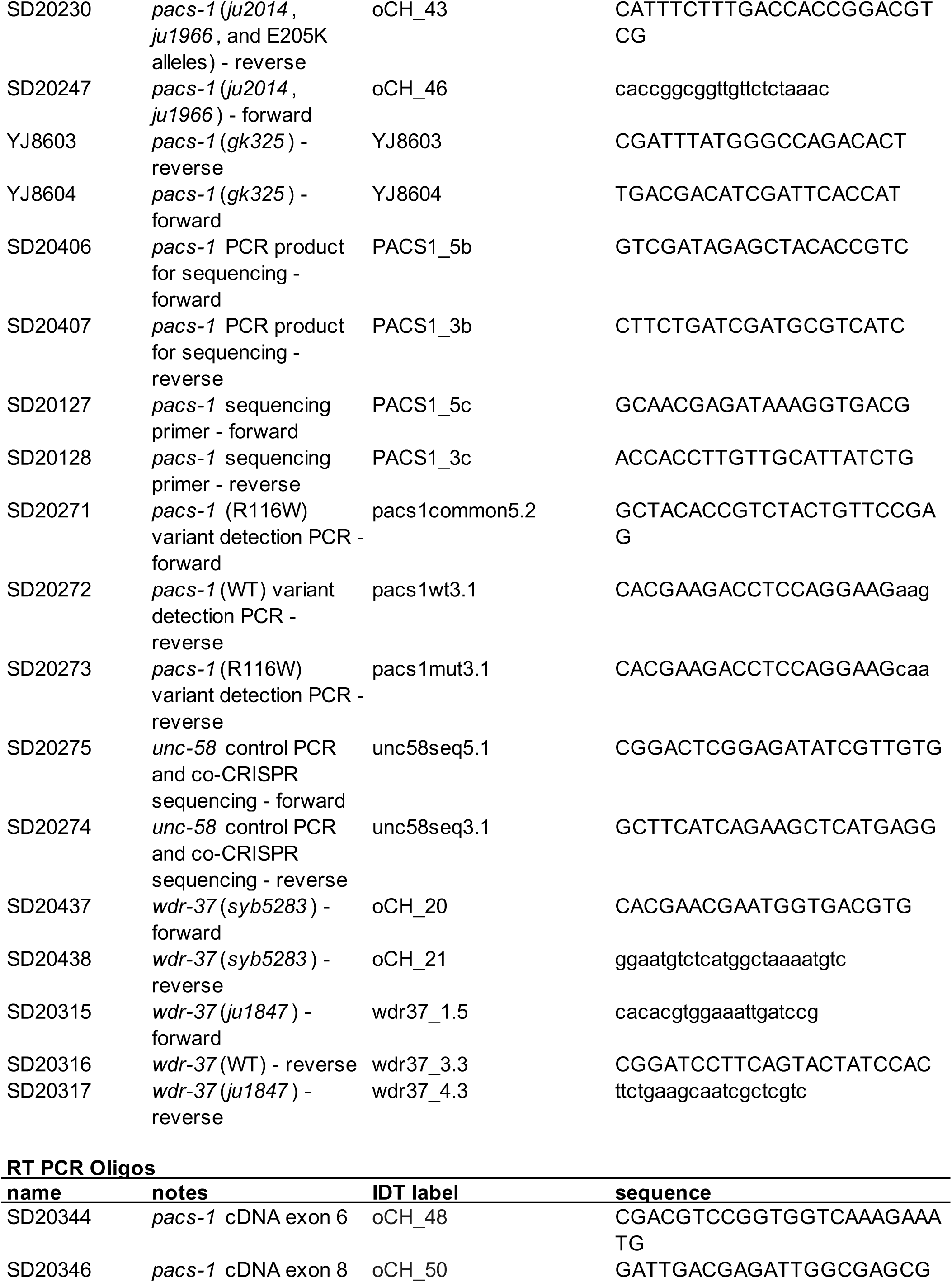
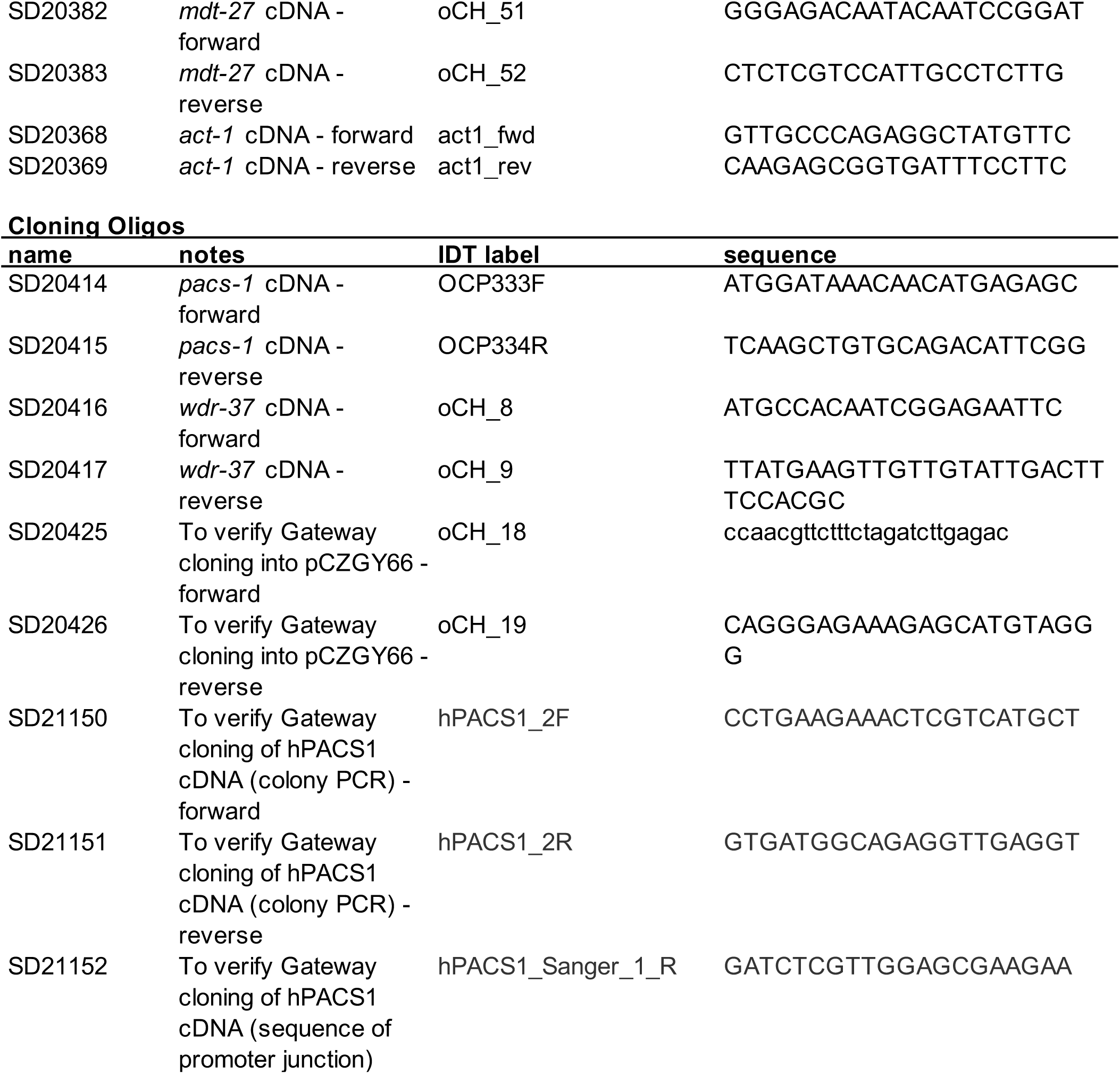
– crRNA and oligo sequences.

**Table S3.**
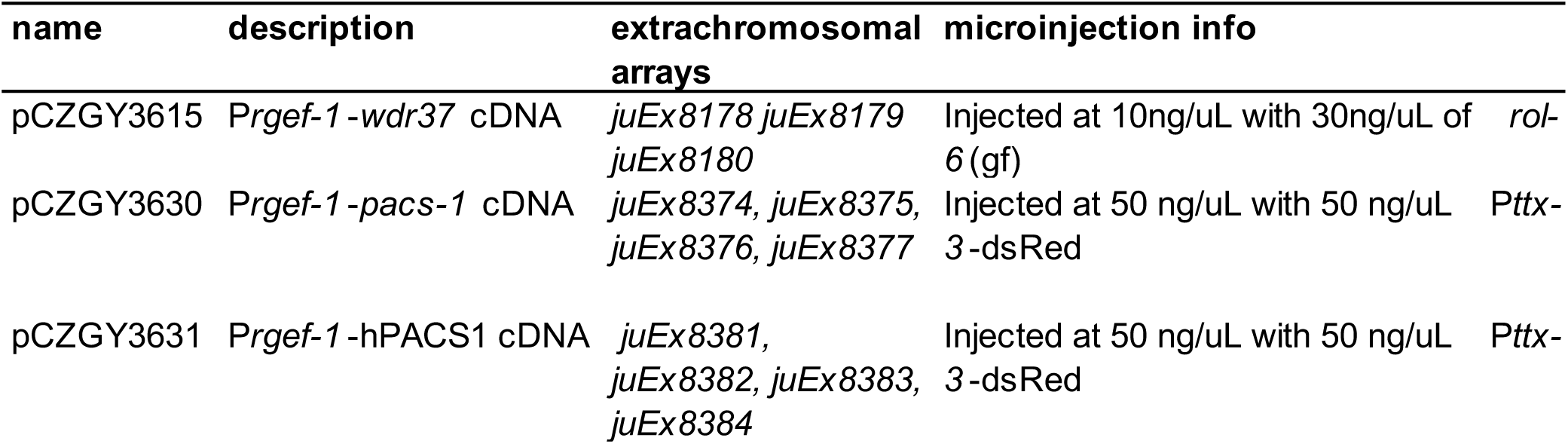
– DNA expression constructs.

## Literature Cited

1. Arribere, J. A., R. T. Bell, B. X. Fu, K. L. Artiles, P. S. Hartman and A. Z. Fire, 2014 Efficient marker-free recovery of custom genetic modifications with CRISPR/Cas9 in Caenorhabditis elegans. Genetics 198: 837–846.

2. Brenner, S., 1974 The genetics of Caenorhabditis elegans. Genetics 77: 71–94.

3. Jenkins, P. M., L. Zhang, G. Thomas and J. R. Martens, 2009 PACS-1 mediates phosphorylation-dependent ciliary trafficking of the cyclic-nucleotide-gated channel in olfactory sensory neurons. J Neurosci 29: 10541–10551.

4. Jospin, M., Y. B. Qi, T. M. Stawicki, T. Boulin, K. R. Schuske et al., 2009 A neuronal acetylcholine receptor regulates the balance of muscle excitation and inhibition in Caenorhabditis elegans. PLoS Biol 7: e1000265.

5. Kanca, O., J. C. Andrews, P. T. Lee, C. Patel, S. R. Braddock et al., 2019 De Novo Variants in WDR37 Are Associated with Epilepsy, Colobomas, Dysmorphism, Developmental Delay, Intellectual Disability, and Cerebellar Hypoplasia. Am J Hum Genet 105: 413–424.

6. Kottgen, M., T. Benzing, T. Simmen, R. Tauber, B. Buchholz et al., 2005 Trafficking of TRPP2 by PACS proteins represents a novel mechanism of ion channel regulation. EMBO J 24: 705–716.

7. Li, S., C. M. Armstrong, N. Bertin, H. Ge, S. Milstein et al., 2004 A map of the interactome network of the metazoan C. elegans. Science 303: 540–543.

8. Liu, H., P. W. Hu, S. Budhiraja, A. Misra, J. Couturier et al., 2020 PACS1 is an HIV-1 cofactor that functions in Rev-mediated nuclear export of viral RNA. Virology 540: 88–96.

9. Malovannaya, A., R. B. Lanz, S. Y. Jung, Y. Bulynko, N. T. Le et al., 2011 Analysis of the human endogenous coregulator complexome. Cell 145: 787–799.

10. Mello, C. C., J. M. Kramer, D. Stinchcomb and V. Ambros, 1991 Efficient gene transfer in C.elegans: extrachromosomal maintenance and integration of transforming sequences. EMBO J 10: 3959–3970.

11. Nair-Gill, E., M. Bonora, X. Zhong, A. Liu, A. Miranda et al., 2021 Calcium flux control by Pacs1-Wdr37 promotes lymphocyte quiescence and lymphoproliferative diseases. EMBO J 40: e104888.

12. Olson, H. E., N. Jean-Marcais, E. Yang, D. Heron, K. Tatton-Brown et al., 2018 A Recurrent De Novo PACS2 Heterozygous Missense Variant Causes Neonatal-Onset Developmental Epileptic Encephalopathy, Facial Dysmorphism, and Cerebellar Dysgenesis. Am J Hum Genet 102: 995–1007.

13. Paix, A., A. Folkmann, D. Rasoloson and G. Seydoux, 2015 High Efficiency, Homology-Directed Genome Editing in Caenorhabditis elegans Using CRISPR-Cas9 Ribonucleoprotein Complexes. Genetics 201: 47–54.

14. Piggott, C. A., Z. Wu, S. Nurrish, S. Xu, J. M. Kaplan et al., 2021 Caenorhabditis elegans junctophilin has tissue-specific functions and regulates neurotransmission with extended-synaptotagmin. Genetics 218.

15. Piguet, V., L. Wan, C. Borel, A. Mangasarian, N. Demaurex et al., 2000 HIV-1 Nef protein binds to the cellular protein PACS-1 to downregulate class I major histocompatibility complexes. Nat Cell Biol 2: 163–167.

16. Reis, L. M., E. A. Sorokina, S. Thompson, S. Muheisen, M. Velinov et al., 2019 De Novo Missense Variants in WDR37 Cause a Severe Multisystemic Syndrome. Am J Hum Genet 105: 425–433.

17. Salzberg, Y., N. J. Ramirez-Suarez and H. E. Bulow, 2014 The proprotein convertase KPC-1/furin controls branching and self-avoidance of sensory dendrites in Caenorhabditis elegans. PLoS Genet 10: e1004657.

18. Schuurs-Hoeijmakers, J. H., E. C. Oh, L. E. Vissers, M. E. Swinkels, C. Gilissen et al., 2012 Recurrent de novo mutations in PACS1 cause defective cranial-neural-crest migration and define a recognizable intellectual-disability syndrome. Am J Hum Genet 91: 1122–1127.

19. Scott, G. K., F. Gu, C. M. Crump, L. Thomas, L. Wan et al., 2003 The phosphorylation state of an autoregulatory domain controls PACS-1-directed protein traffic. EMBO J 22: 6234–6244.

20. Sieburth, D., Q. Ch’ng, M. Dybbs, M. Tavazoie, S. Kennedy et al., 2005 Systematic analysis of genes required for synapse structure and function. Nature 436: 510–517.

21. Sorokina, E. A., L. M. Reis, S. Thompson, K. Agre, D. Babovic-Vuksanovic et al., 2021 WDR37 syndrome: identification of a distinct new cluster of disease-associated variants and functional analyses of mutant proteins. Hum Genet 140: 1775–1789.

22. Thomas, G., J. E. Aslan, L. Thomas, P. Shinde, U. Shinde and T. Simmen, 2017 Caught in the act – protein adaptation and the expanding roles of the PACS proteins in tissue homeostasis and disease. J Cell Sci 130: 1865–1876.

23. Waites, C. L., A. Mehta, P. K. Tan, G. Thomas, R. H. Edwards and D. E. Krantz, 2001 An acidic motif retains vesicular monoamine transporter 2 on large dense core vesicles. J Cell Biol 152: 1159–1168.

24. Wan, L., S. S. Molloy, L. Thomas, G. Liu, Y. Xiang et al., 1998 PACS-1 defines a novel gene family of cytosolic sorting proteins required for trans-Golgi network localization. Cell 94: 205–216.

25. Youker, R. T., U. Shinde, R. Day and G. Thomas, 2009 At the crossroads of homoeostasis and disease: roles of the PACS proteins in membrane traffic and apoptosis. Biochem J 421: 1–15.

26. Zang, R. X., M. J. Mumby and J. D. Dikeakos, 2022 The Phosphofurin Acidic Cluster Sorting Protein 2 (PACS-2) E209K Mutation Responsible for PACS-2 Syndrome Increases Susceptibility to Apoptosis. ACS Omega 7: 34378–34388.

